# A two-type branching process model of gene family evolution

**DOI:** 10.1101/2021.03.18.435925

**Authors:** Arthur Zwaenepoel, Yves Van de Peer

**Affiliations:** Department of Plant Biotechnology and Bioinformatics, Ghent University, 9054 Ghent, Belgium; Center for Plant Systems Biology, VIB, 9054 Ghent, Belgium; Department of Biochemistry, Genetics and Microbiology, University of Pretoria, Pretoria 0028, South Africa; College of Horticulture, Nanjing Agricultural University, Nanjing, 210095, China

**Keywords:** gene duplication and loss, phylogenetics, multi-type branching process, Bayesian inference

## Abstract

Phylogenetic models of gene family evolution based on birth-death processes (BDPs) provide an awkward fit to comparative genomic data sets. A central assumption of these models is the constant per-gene loss rate in any particular family. Because of the possibility of partial functional redundancy among gene family members, gene loss dynamics are however likely to be dependent on the number of genes in a family, and different variations of commonly employed BDP models indeed suggest this is the case. We propose a simple two-type branching process model to better approximate the stochastic evolution of gene families by gene duplication and loss and perform Bayesian statistical inference of model parameters in a phylogenetic context. We evaluate the statistical methods using simulated data sets and apply the model to gene family data for *Drosophila*, yeasts and primates, providing new quantitative insights in the long-term maintenance of duplicated genes.

## Introduction

The expansion and contraction of gene families by gene duplication and loss is one of the key features of genome evolution. Statistical phylogenetic analyses of the evolution of gene family sizes rely on probabilistic models of gene family evolution, which are typically based on birth-death processes (BDPs) (Hahn et al. 2005; Csűrös and Miklós 2009). Phylogenetic BDP models have been applied to analyze genome evolution using comparative genomic data sets (e.g. Hahn, Han, and Han 2007; Csűrös and Miklós 2009; Thomas et al. 2020), infer reconciled gene trees (Åkerborg et al. 2009; Szöllősi et al. 2015), and infer ancient whole-genome duplications in a phylogeny (Rabier, Ta, and Ané 2014; Zwaenepoel and Van de Peer 2020). Importantly, such models seek to capture the long-term stochastic evolution of gene family sizes by duplication and loss, as opposed to the short-term population genetic processes leading to the establishment and loss of newly arisen copy number variants.

Standard phylogenetic BDP models provide however an awkward fit to comparative genomic data. Our study departs from the observation that estimated gene loss rates can vary dramatically depending on the data and assumed BDP model. For instance, when considering the full array of gene families across a set of eight *Drosophila* species, the standard linear BDP model yields an estimated loss rate of 0.24 expected loss events per gene per 100 My; whereas considering the set of gene families that are retained with at least one copy across all taxa, and assuming these gene families cannot go extinct (i.e. are essential), a loss rate of 3.94 expected loss events per *duplicated* gene per 100 My is obtained (see below for methods). The same discrepancy appears when comparing loss rate estimates from the linear BDP model with crude estimates based on age distributions of duplicated genes (Lynch and Conery 2003).

One of the key assumptions underlying the standard linear BDP model that is generally violated, and which may lead to the discrepancies noted above, is the assumption of a single and constant loss rate per gene in a family. Many gene duplicates, although stably established in the genome, exhibit some functional redundancy (examples abound in the molecular biological literature). When a duplicated gene is fully or partially functionally redundant with its parental gene, it seems likely that both copies are subject to higher gene loss rates until either one gene of the duplicate pair is lost or has adopted a distinct function (neofunctionalization), or both genes underwent ‘complete’ subfunctionalization (hereafter, neo- and subfunctionalization are referred to jointly as ‘x-functionalization’). In other words, when a set of genes is (partially) functionally redundant, we may expect an increased per-gene loss rate *µ*_*r*_ compared to a set of non-redundant genes with per-gene loss rate *µ*_*nr*_, as in the former case there will be weaker purifying selection against pseudogenes (in the case of full redundancy for instance, a null-mutant will be effectively neutral (Walsh 2003)). This however induces non-independence among distinct genes in a family, because when one copy in a functionally redundant gene pair gets lost, the loss rate of the retained copy will drop back to *µ*_*nr*_. This contrasts with the commonly employed BDP models which obey the branching property that distinct genes evolve independently. This non-independence represents a serious obstacle for developing more accurate stochastic models of gene family evolution.

In this study, we propose a two-type continuous-time branching process model of gene family evolution to better approximate the stochastic evolution of gene families, while retaining the independence assumption. In this model we treat a family as consisting of two types of genes: ‘base’ genes (type 1) and ‘excess’ genes (type 2). Any particular family at time *t* can be represented as a tuple *X*(*t*) = (*X*_1_(*t*), *X*_2_(*t*)) where *X*_1_(*t*) and *X*_2_(*t*) represent the number of type 1 and type 2 genes in the family at time *t* respectively. We can think of such a family as consisting of *Z*(*t*) = *X*_1_(*t*) + *X*_2_(*t*) genes, coding for *X*_1_(*t*) functions (note that we define a gene family as an *orthogroup*, irrespective of gene function, see methods). We allow for four types of events: gene duplication at rate *λ* per gene, loss of a type 1 gene at rate *µ*_1_ per type 1 gene, loss of a type 2 gene at rate *µ*_2_ per type 2 gene and a conversion of a type 2 to a type 1 gene (x-functionalization) at a rate *ν* per type 2 gene (see methods).

Importantly, because of the independence assumption, *µ*_2_ cannot be directly interpreted as the rate of gene loss of a functionally redundant gene *µ*_*r*_. To see this, consider a state, say, *X*(*t*) = (1, 3), which we could interpret as a set of four genes coding for a single function. All four genes are functionally redundant in this case, and under the ideal model we would have a total loss rate 4*µ*_*r*_ where *µ*_*r*_ is the loss rate for a functionally redundant gene. Instead under the two-type model, the total rate of gene loss will be *µ*_1_ + 3*µ*_2_, so that when *µ*_1_ ≪ *µ*_2_ the rate of loss per redundant gene is ≈ 3*µ*_2_/4. Under the two-type model the rate of gene loss per *excess* gene *µ*_2_ is constant for different family sizes, while the rate of gene loss per *redundant* gene increases with increasing number of excess genes per base gene, approximately equaling *µ*_2_(*n* − 1)/*n* for a group of *n* redundant genes. Another example illustrating the differences between the ideal model with non-independence and the two-type model is provided in fig. 1 (a).

**Figure 1:**
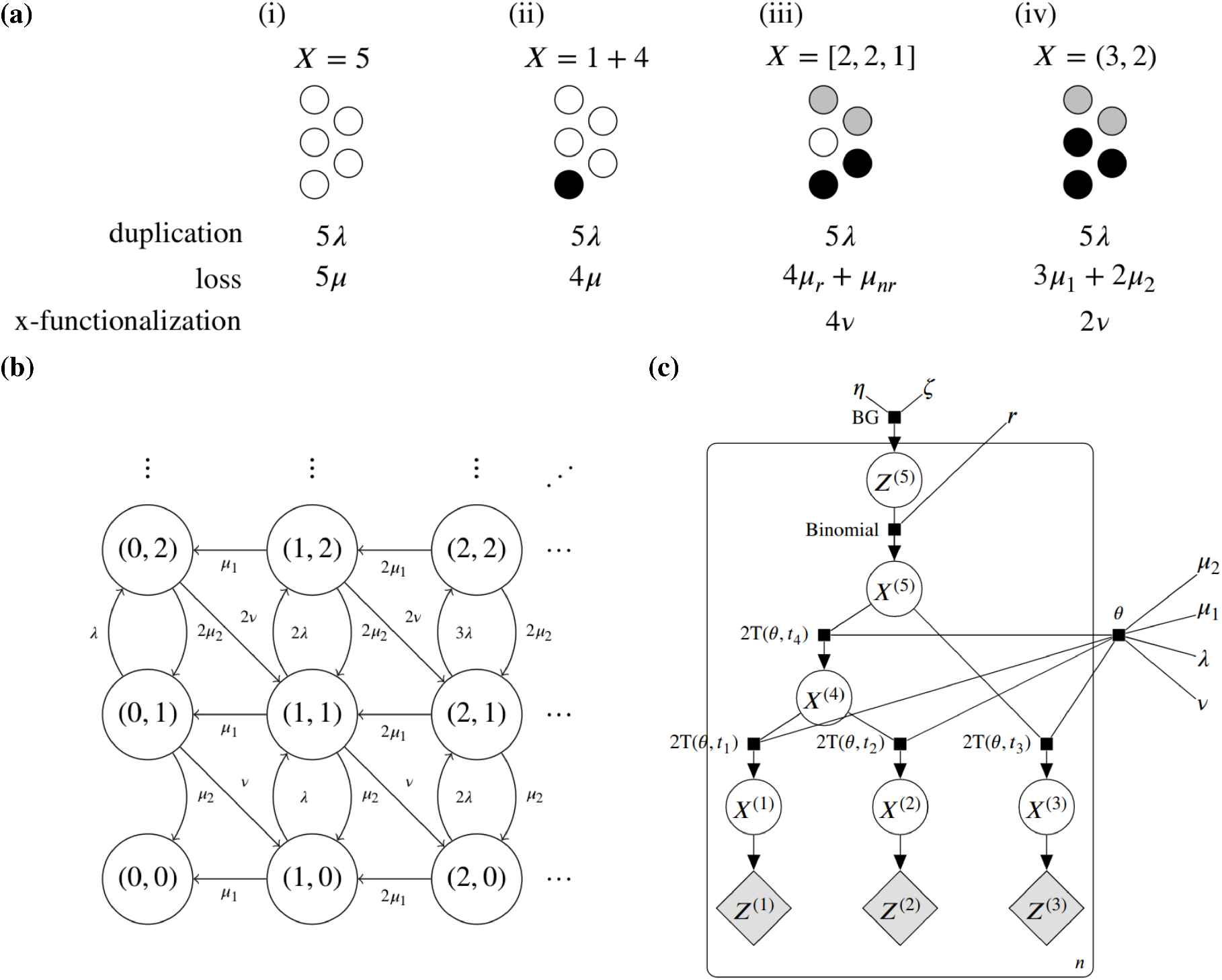
(a) Schematic representation of a gene family of total size *Z* = 5, consisting of a set of genes coding for three functions, with its associated state *X* from the perspective of four models considered in this paper. The associated total (i.e. family-level) duplication, loss and x-functionalization rates are indicated below. In all models the per-gene duplication rate is *λ*. (i) Simple linear BDP model with per-gene loss rate *µ*; (ii) Linear BDP without extinction, with loss rate per *excess* gene (in white) *µ*; (iii) Duplication-loss-functionalization (DLF) model. The five gene copies are explicitly subdivided in three functional groups (depicted in different colors). Genes in groups consisting of multiple copies (gray and black genes) are subject to loss at rate *µ*_*r*_ per gene, while genes in single-copy groups are subject to loss at rate *µ*_*nr*_ < *µ*_*r*_. Genes in multi-copy groups give rise to new groups (x-functionalize) at rate *v* per gene; (iv) Two-type duplication-loss model with loss rate *µ*_1_ per *base* (type 1, in black) gene, loss rate *µ*_2_ per *excess* (type 2, in gray) gene and x-functionalization rate *v* per type 2 gene. (b) Graph representation of the two-type continuous time branching process, where nodes depict states *X* = (*X*_1_, *X*_2_) and edges transitions occuring with the associated rates. (c) Simplified probabilistic graphical model for the two-type DL model along a three-taxon phylogeny, following the directed factor graph notation of Dietz (2010). BG denotes the Beta-Geometric distribution and 2T(*θ, t*_*u*_) denotes the transient distribution for the two-type branching process with parameters *θ* along the branch leading to node *u* of length *t*_*u*_. Priors for *θ* = (*λ, µ*_1_, *v, µ*_2_) and hyperparameters *ϕ* = (*η, ζ, r*) are not shown.

In the remainder of this paper we describe the probabilistic model and an approach for Bayesian statistical inference of the relevant model parameters from comparative genomic data. We also present some innovations that apply more generally to phylogenetic BDP models, in particular the usage of a Beta-Geometric prior on the number of genes at the root and a BDP model for families that are assumed not to go extinct. We study the statistical properties of inference under the two-type branching process model using simulated data sets and study gene family evolution in *Drosophila*, yeasts and primates. Finally, we discuss the implications of our results for the long-term evolutionary dynamics of gene content and the evolutionary genetics of gene duplication.

## Model and methods

### Duplication-loss-functionalization model

We first describe an idealized duplication-loss-functionalization (DLF) model with non-independent evolution of gene copies. Throughout, we define a gene family as an orthogroup: a group of genes across a set of species derived from a common ancestral gene in the last common ancestor of the species under consideration. We assume a gene family of *n* genes is partitioned in *k* ≤*n* functions, and we represent a gene family at time *t* as the length *k* array of gene counts in each functional class *X*(*t*) = [*X*_1_(*t*), *X*_2_(*t*), …, *X*_*k*_(*t*)], *X*_*i*_(*t*) > 0, *I* ≤ *k* An extinct gene family is represented by the empty array. The total gene family size is denoted by 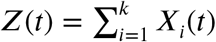 An example gene family under this model with *Z* = 5 and *k* = 3 is shown in fig. 1 (a.iii). We assume birth-death dynamics, i.e. transitions increment or decrement *Z*(*t*) by at most one unit. We assume all genes duplicate at rate *λ* and that genes in a functional class *i* with *X*_*i*_ > 1 suffer loss at rate *µ*_*r*_ per gene (loss of a redundant copy). Genes in a single-copy functional class suffer loss at rate *µ*_*nr*_ (loss of a non-redundant copy), an event that leads to a decrease in the number of functional classes in the family. Lastly, we assume that for each class *i* for which *X*_*i*_ > 1, genes shift to a new, not yet existing functional class at rate *v* per gene, decrementing *X*_*i*_ and increasing the number of functional classes in the family (neo- or subfunctionalization). Note that the latter event does not affect *Z*(*t*). While this model does not admit efficient statistical inference (as far as we are aware), it is straightforward to simulate from using standard techniques.

### Two-type branching process model

As an approximation to the idealized DLF model, we model the evolution of gene family content along a species tree as a two-type homogeneous continuous-time branching process *X*(*t*) = (*X*_1_(*t*), *X*_2_(*t*)) ∈ ℕ×ℕ (where we take ℕ to include 0). Here *X*_1_(*t*) and *X*_2_(*t*) denote the number of type 1 and type 2 genes at time *t* respectively (fig. 1 a.iv). The stochastic evolution of the process is determined by the rate parameters *θ* = (*λ, µ*_1_, *µ*_2_, *ν*) in the following way

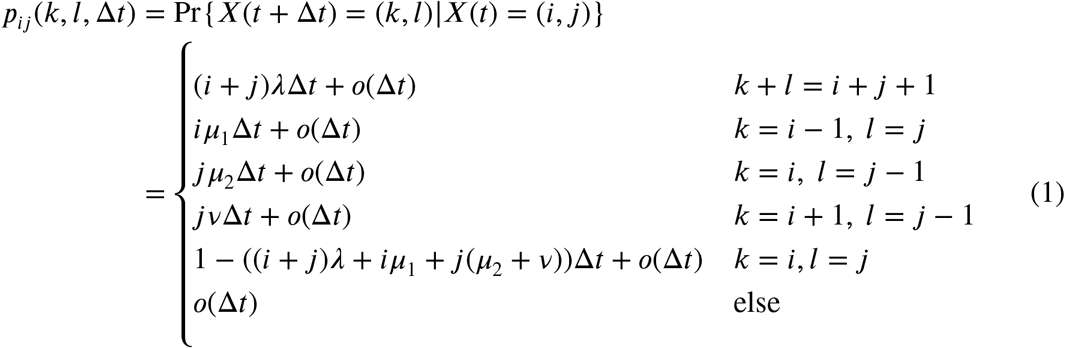

for Δ*t* sufficiently small and *i, j, k, l* ≥ 0. The process is a bivariate birth-death process and is Markovian (see fig. 1 (b) for a graph representation of the bivariate Markov chain). Throughout we assume {*µ*_1_, *λ, v*} < *µ*_2_. As a model of gene family evolution by gene duplication and loss, *X*_1_(*t*) denotes the number of ‘base’ genes in the family, while *X*_2_(*t*) denotes the number of cess’ genes in the family for which the eventual fate (non, neo-subfunctionalization) is yet be determined at time *t*. The model defined above assumes that genes duplicate at the same per-gene rate *λ* and that the fate of a type 2 gene is resolved after an exponentially distributed time with mean *µ*_2_ + *v*, with the probability of nonfunctionalization being *µ*_2_/(*µ*_2_ + *v*) and the probability of sub- or neofunctionalization *v*/(*µ*_2_ + *v*) (turning an excess gene into a base gene). We further assume type 1 genes are removed with rate *µ*_1_. We refer to this model as the ‘two-type duplication-loss (DL) model’.

### Transition probabilities for the two-type model

Probabilistic inference of model parameters from comparative genomic data requires that we can compute transition probabilities under the model. In general, the transient distributions for multi-type branching processes like the above are analytically intractable, presenting a serious obstacle to their fruitful application in a statistical setting. Xu et al. (2015) however presented a numerical approach for computing transition probabilities for Markovian multi-type branching processes based on inversion of the associated probability generating functions (pgfs). Let *f*_*ij*_(*s*_1_, *s*_2_, *t*) denote the pgf for the two-type DL model in eq. 1 when starting at *X*(0) = (*i, j*), i.e.

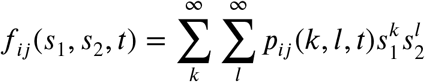

Importantly, the branching property implies the following relationship among the probability generating functions for different *X*(0)

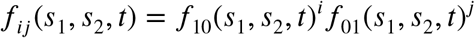

(e.g. Athreya and Ney (1972)), so we may work with *f*_10_ and *f*_01_ and recover the desired pgfs easily. As we derive in detail in Appendix 1, we can find the following system of ordinary differential equations (ODEs) for the pgfs

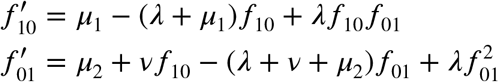

Where the arguments *s*_1_, *s*_2_ and *t* are omitted for clarity and differentiation is with respect to *t*.

No closed form solution for *f*_10_(*s*_1_, *s*_2_, *t*) and *f*_01_(*s*_1_, *s*_2_, *t*) can be obtained for this system, and we solve these ODEs numerically using the TSit5 solver implemented in DifferentialEquations.jl (Rackauckas and Nie 2017; Tsitouras, Famelis, and Simos 2011). To obtain transition probabilities from the pgfs we use the numerical inversion method of Xu et al. (2015), which involves re-expressing the pgf as a Fourier series 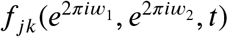 so that the coefficients corresponding to the transition probabilities are given by the inverse Fourier transform, which can be computed numerically using the fast Fourier transform (FFT) along an *N* × *N* grid. Choice of a higher *N* should lead to more accurate transition probabilities, and determines the maximum state from and to which we can compute transition probabilities. In practice the numerical error is dominated by the tolerance settings in the ODE solver, so that increasing *N* beyond relatively small values (e.g. *N* = 16 as in Xu et al. (2015)) does not lead to much gain in accuracy for transition probabilities among states with reasonable probability in our applications (fig. S1).

### Count data likelihood along a phylogeny

With a reasonably efficient method for computing the transition probabilities available, likelihood-based statistical inference is possible. Our data set is an *n* × *m* matrix consisting of gene counts for *n* gene families for a set of *m* species related by a known phylogeny *S*. We refer to each row *y*_*i*_, *i* ∈ (1, …, *n*), of this matrix as a *phylogenetic profile*. In our notation, the profiles correspond to the *m*-tuples 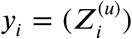 with *u* ∈ {1, …, *m*}, where 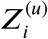 denotes the *total* gene count at leaf node *u* for family *i*, such that 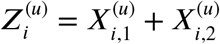 (where 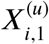 denotes the number of type 1 genes at leaf node *u* for family *i*).

The model we consider assumes that the two-type branching process operates along the branches of species tree *S*, such that at each split in the tree, two independent copies of the ancestral process start with the same initial state. We assume phylogenetic profiles are independent and identically distributed (iid) under the two-type branching process model defined along *S*, and that we have no access to any information on the number of genes of each type at leaf nodes (i.e. we only observe *Z*, not *X*). The resulting probabilistic graphical model (PGM) is depicted for a hypothetical three-taxon tree (*m* = 3) in fig. 1. The likelihood of the observed data 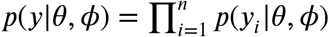 conditional on the parameters of the branching process *θ* and the prior distribution^*i*=^f^1^or the root *ϕ* can be computed using variable elimination along the PGM (i.e. using Felsenstein’s pruning algorithm, and integrating the marginal likelihood values at the root over a suitable prior distribution on the root state).

### Conditioning on non-extinction

To rule out *de novo* gain of genes in arbitrary subtrees of the phylogeny, we filter the data so that at least one gene is present in each clade stemming from the root of the species tree (as in e.g. Rabier, Ta, and Ané (2014) and Zwaenepoel and Van de Peer (2019)). We denote the event of extinction below a node *u* in the species tree by *E*_*u*_ and let *a, b* and *c* respectively label the root and its two daughter nodes. The likelihood of a profile *y* conditional on the event of non extinction in both clades is then

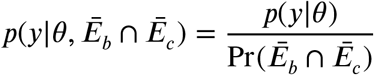

where *Ē* denotes the negation (or complement) of *Ē*. To obtain the relevant data likelihood we therefore need to compute the condition factor Pr(*Ē* _*b*_ n *Ē* _*c*_). Assuming a suitable prior distribution on the number of lineages of each type at the root of the species tree (see below), we can rely on the conditional independence of subtrees of *S* and properties of the pgfs to obtain this probability. For ease of notation, denote by 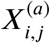 the event that *X*^(*a*)^ = (*i, j*). The condition factor is then given by

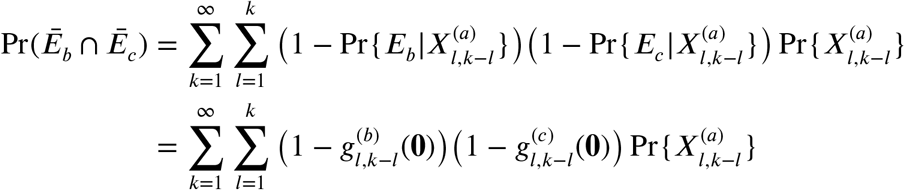

Here 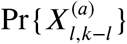 is given by the prior distribution on the number of ancestral lineages in a gene family at the root of the species tree, and 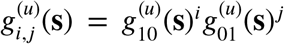 is the joint pgf for the leaf observations in the subtree rooted in node *u*, conditional on the state at the parent of *u*, say *u*’ being (*i, j*). Assuming there are *m* leaves below *u* and labeling the entries of **s** as (*s*_1,1_, *s*_1,2_, *s*_2,1_, *s*_2,2_, …, *s*_*m*,1_, *s*_*m*,2_), where the first index refers to the leaf node in the subtree below *u* and the second to the gene type, this pgf is written more explicitly as

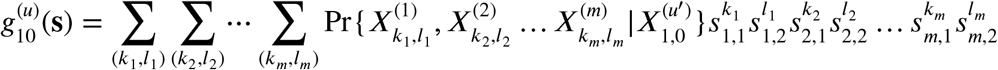

and of course analogously for 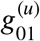. By the branching property and conditional independence of disjoint subtrees, this pgf can be evaluated efficiently using a postorder traversal and the single branch pgf *f*_10_. Specifically, consider a node *u*, a distance *t*_*u*_ from its parent, with (if it is not a leaf) child nodes *ν* and *w*. We have the following recursive relation:

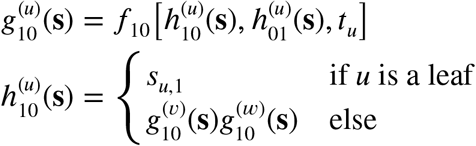

with analogous recursions holding for 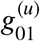 and 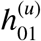.

### Prior distribution on the number of lineages at the root in a gene family

Plausible inferences for the proposed model do not only depend on the stochastic process describing the transient evolution of gene family counts, but also on the assumptions of the ancestral gene content for a family. Since neither the single-type linear BDP nor the two-type DL model have a stationary distribution on ℕ, the model itself does not directly suggest a natural prior for the number of ancestral lineages. Conditional on non-extinction however, the linear birth-death process has a Geometric(*η*) stationary distribution Pr{*X* = *k*} = *η*(1 −*η*)^*k*−1^, and a Geometric prior has been considered previously in this context (Rabier, Ta, and Ané 2014; Zwaenepoel and Van de Peer 2019).

The size distribution of gene families is however universally overdispersed with respect to the geometric distribution, showing an approximate power-law tail (Huynen and Van Nimwegen 1998). As we plan to study in detail elsewhere, the major cause of the apparent power-law frequency distribution is likely heterogeneity in gene duplication and loss rates across families, a hypothesis also investigated by Hughes and Liberles (2008). A simple model showing this behavior, corresponding to a special case of the two-type process with *µ*_1_ = *v* = 0, is derived in Appendix 2. The resulting stationary distribution is a Beta-Geometric BG(*α, β*) distribution, which has probability mass function (pmf)

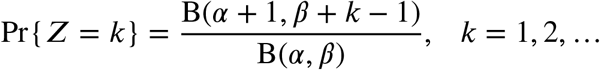

Where B(*α, β*) denotes the beta function 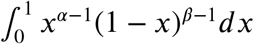.This pmf shows approximate power-law behavior in its tail when *α* and *β* are fairly small. The BG distribution provides an excellent fit to the observed size distribution for non-zero gene counts (fig. 2). Note furthermore that the mean of the underlying beta distribution (see Appendix 2) may provide a crude idea of the ratio *λ/µ*_2_ in the two-type model.

**Figure 2:**
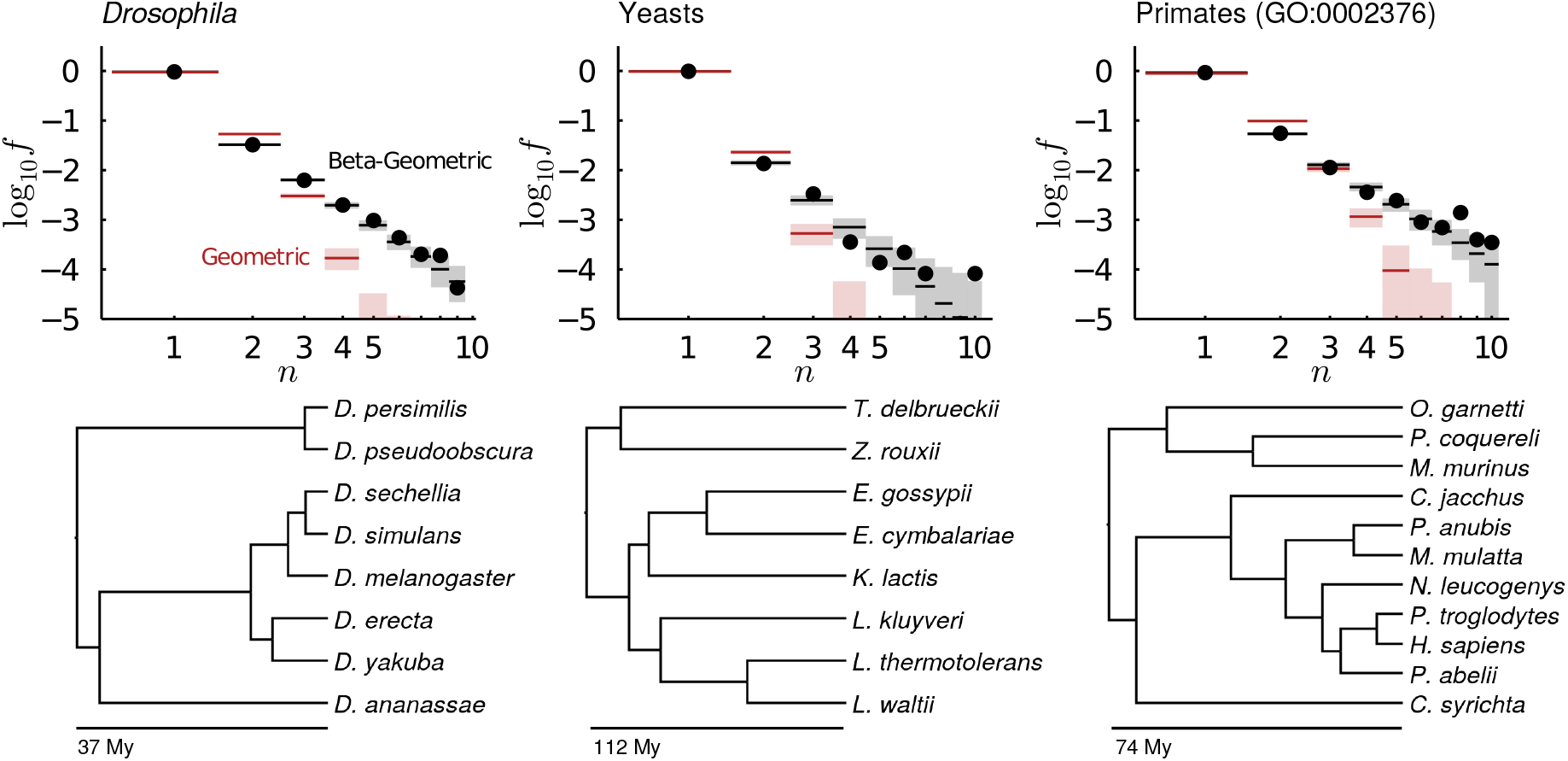
Top row: Geometric and Beta-Geometric stationary distributions fitted to observed gene family size distributions for three data sets. The black dots show the observed size distribution (across all taxa), whereas the lines show the mean frequencies and 95% posterior predictive intervals based on 1000 simulations from the posterior predictive distribution for both models. Bottom row: Phylogenies for the relevant data sets, refer to table S1 for complete taxon names.

The BG distribution is a reasonable prior distribution for the *total* number of genes *Z*^(*a*)^ at the root node *a* of the species tree, however for inference purposes a prior for *X*^(*a*)^ is required. Here we assume that there is at least one type 1 gene in each family, and that among the *Z*^(*a*)^ − 1 remaining genes, each gene is of type 1 with probability *r* and type 2 with probability 1 − *r*, so that the number of type 2 genes is a Binomial(*Z*^(*a*)^ − 1, 1 − *r*) random variable. We use the resulting distribution for *X*^(*a*)^ in our analyses for the two-type DL model (fig. 1) and refer to it as the BG-Binomial prior. Importantly, while we motivate the usage of the BG distribution using the fact that the observed size distribution is overdispersed with respect to the geometric distribution (and that this is likely caused by rate heterogeneity across families, see Appendix 2), we do ignore this rate heterogeneity in the transient regime for reasons of computational efficiency (but see fig. 5 and associated discussion).

**Figure 5:**
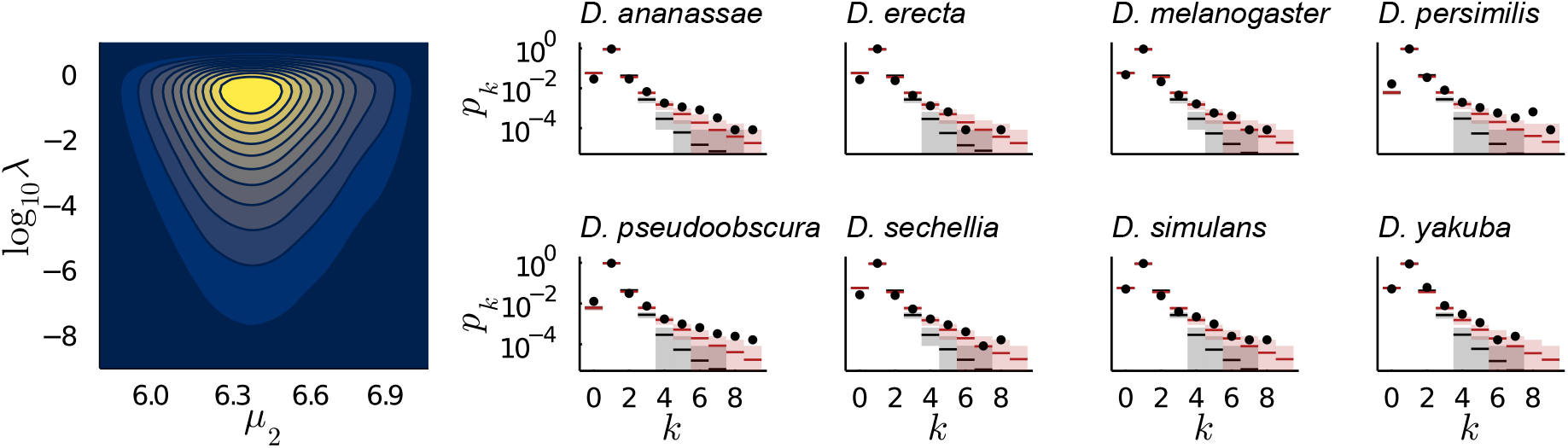
Posterior predictive simulations assuming *λ*/*µ*_2_ is Beta distributed across gene families with dispersion *ζ* = 4.01. On the left the joint distribution of family-specific *λ* and posterior mean *µ*_2_ values is shown. The eight panels on the right show the posterior predictive family size distributions with (red) and without (black) the *ad hoc* procedure to account for rate heterogeneity across families based on the Beta-Geometric assumption.

### Bayesian inference and posterior predictive simulation

With an algorithm for the likelihood (*p y*|*θ, ϕ)* available we can perform Bayesian inference for the two-type DL model using Markov Chain Monte Carlo (MCMC) given suitable prior distributions. Unless stated otherwise, we adopt the following priors in our analyses:

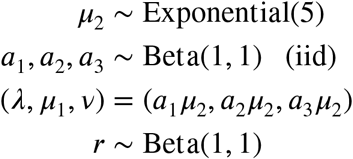

Assuming {*λ, µ*_1_, *v*} < *µ*_2_. Unless stated otherwise, we fix *η* and *ζ* to the posterior mean values obtained from the fit of the stationary distribution of the single-type model without extinction to the relevant data (see fig. 2). We implemented a simple adaptive Metropolis-within-Gibbs (MWG) algorithm (Roberts and Rosenthal 2009) to sample from the posterior distribution. We make extensive use of posterior predictive simulations to assess model fit. Throughout, posterior predictive simulations are conducted by sampling 1000 random parameter vectors from the joint posterior distribution. For each sample we simulate a data set of equal size as the data set for which inference was performed, and for each such simulation replicate we compute the size distribution for each lineage and across all lineages.

### Single-type BDP models

We compare analyses under the two-type DL model with analyses for three distinct single-type BDP models. First, we consider a critical linear BDP model where genes duplicate and get lost at a single ‘turnover’ rate *λ*. As a second model, we consider the default linear BDP with duplication and loss rates *λ* and *µ* per gene respectively. Lastly, we consider the single-type linear BDP model without extinction (see Appendix 2), where genes duplicate at rate *λ*, and the loss rate per *excess* gene is *µ*. The latter model is equivalent to a linear birth-death and immigration process (or duplication, loss and gain model in Csűrös and Miklós (2009)) on the number of excess genes (i.e. *Z* − 1 instead of the family size *Z*) with immigration (gain) rate, *κ* = *λ*. We note that a discrete-time deterministic analog of this model was used by Lynch and Conery (2003) to model age distributions of duplicated genes. An illustration of how these various models differ in terms of family-level duplication and loss rates from each other, the DLF model and the two-type DL model is provided in fig. 1 (a). The single-type model without extinction can further be seen as a special case of the two-type DL model where *µ*_1_ = *v* = 0. Inference for single-type models is performed using the DeadBird.jl library (which builds on Zwaenepoel and Van de Peer 2020), which employs the algorithm for computing the phylogenetic BDP likelihood devised by Csűrös and Miklós (2009). We further make use of the Turing.jl library (Ge, Xu, and Ghahramani 2018) using the NUTS algorithm (Hoffman and Gelman 2014) for Bayesian inference. We use the same BG prior as for the two-type model as a distribution on the number of lineages at the root (see Appendix 3) and use non-informative (flat) priors for the rate parameters unless stated otherwise. Posterior predictive simulations were performed as for the two-type DL model.

### Data sets, implementation and availability

We consider three data sets, consisting of (1) eight *Drosophila* species (a subset from the widely studied 12 *Drosophila* data set (Hahn, Han, and Han 2007; Drosophila 12 Genomes Consortium 2007)), (2) eight yeast species (the set of species included in the yeast gene order browser (YGOB, v7-Aug2012; Byrne and Wolfe (2005)) that did not undergo the *S. cerivisiae* genome duplication) and (3) 11 primate species. Gene families for each data set were obtained using OrthoFinder (v2.4) (Emms and Kelly 2019), using the sequence data listed in table S1 (when the data included isoforms, we retained the longest isoform as a representative for each gene). For the sake of computational feasibility, we focus for the primates phylogeny on a subset of the data by identifying those families that contain a human homolog annotated with the gene ontology term GO:0002376 (immune system process) or any of its descendant terms. The data was filtered such that there was at least one representative for each family in both clades stemming from the root. For computational feasibility, we further excluded families with more than 10 copies in any of the included taxa, so that in the end we obtain a data set containing 12.163, 4855 and 1921 gene families for the *Drosophila*, yeast and primates phylogenies respectively. For the *Drosophila* and primates data sets, a dated phylogeny was downloaded from the TimeTree database (Kumar et al. 2017) (fig. 2). For the yeast data set, we inferred a dated time tree using r8s (Sanderson 2003) under the molecular clock assumption, using the phylogeny inferred by OrthoFinder and a root calibration of 112 My in line with Beimforde et al. (2014). All methods are implemented in the Julia programming language (Bezanson et al. 2017) and the associated packages are freely available at https://github.com/arzwa/TwoTypeDLModel and https://github.com/arzwa/DeadBird.jl. The data sets and scripts for conducting the analyses performed in this paper are also available from the former repository.

## Results

### Parameter estimation for simulated data under the two-type DL model

We first assess our ability to recover true parameter values of the two-type DL model for simulated data from the same model. In particular, the fact that in real empirical data sets we can only observe the total gene count *Z*^(*u*)^ for each family at each leaf node *u* (and not the counts for each type 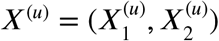) is a potential source of identifiability issues. Indeed, it is clear that for a single branch, it would be impossible to identify model parameters of the two-type DL model based on nothing more than a single observation of *Z* for each family. By considering gene family counts along a phylogeny, a single family provides multiple correlated observations of the evolutionary process, providing information about gene content in ancestral branches of the tree. Observations of *Z* along the leaves of a phylogeny should therefore also provide information about ancestral gene content at the type level (*X*_1_, *X*_2_).

Simulations of data sets along the *Drosophila* phylogeny consisting of 1000 gene families across a range of randomly drawn parameter values indicate that overall, parameters can be estimated accurately even for relatively small data sets (fig. S2). As expected, using the completely observed two-type data does lead to a considerably lower variance in the marginal posterior distribution for *v* compared to the collapsed data. It may be that the marginal posterior mean for *v* slightly overestimates the true value when using the incompletely observed data, however all uncertainty intervals contain the true value, so this risk seems to be minor. Also noticeable, but less dramatic is the difference in posterior variance for *µ*_2_ and *r* (the probability of excess genes at the root to be of type 1). In addition, we simulated a data set of 10.000 gene families along the eight-taxon *Drosophila* phylogeny using model parameters that seemed reasonable based on exploratory analyses of subsets of the actual *Drosophila* data set (*λ* = 0.2, *µ*_1_ = 0.1, *v* = 0.2, *µ*_2_ = 5, *η* = 0.95, *ζ* = 4 and *r* = 0.5). We obtain similar results as for our smaller simulations (fig. 3), in particular we find that the posterior variance for *v* is much higher when using the incompletely obeserved data. Nevertheless, the associated posterior mean values seem to align rather well with each other and the true value, suggesting that inference for the two-type model in a phylogenetic context is indeed possible without observing a type-specific census. Very similar observations can be made for independent replicate simulations (fig. S3, fig. S4). When we perform parameter inference for the default single-type BDP model and the single-type model without extinction for the same simulated data set, we find that the duplication rate tends to be underestimated in both models (table S2). The loss rate of the default single-type model only very slightly overestimates *µ*_1_ while the loss rate in the single-type model without extinction strongly underestimates *µ*_2_.

**Figure 3:**
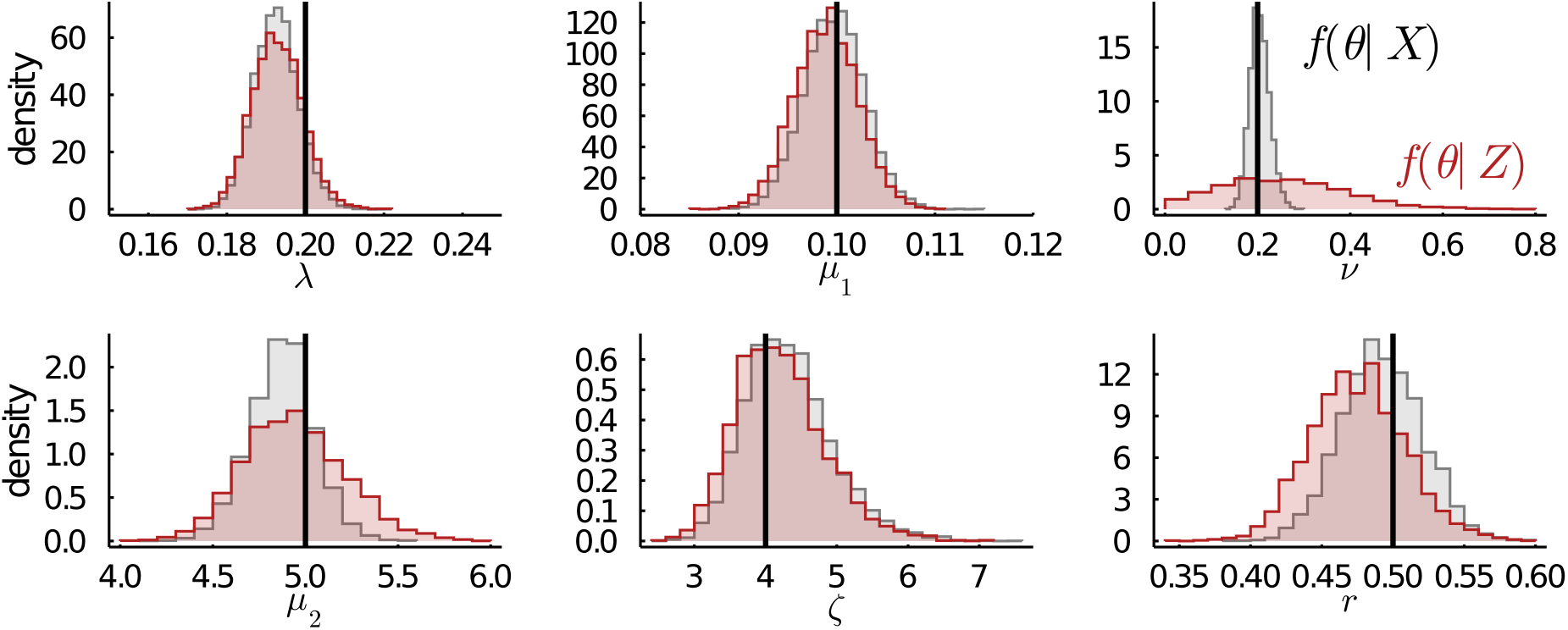
Marginal posterior distributions for a simulated data set of 10.000 gene families, simulated along the eight-taxon *Drosophila* phylogeny from the two-type DL process with parameters *λ* = 0.2, *µ*_1_ = 0.1, *ν* = 0.2, *µ*_2_ = 5 and a bounded Beta-geometric prior on the number of lineages at the root with *η* = 0.95, dispersion *ζ* = 4 and bound at *Z* = 10. When there is more than one lineage at the root, each additional lineage has a *r* = 0.5 probability of being a type 2 gene. In black the posteriors conditional on the fully observed data (i.e. 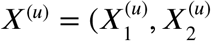) tuples for each leaf node *u*) are shown, while in red the posteriors conditional on the total gene count *Z*^(*u*)^ are shown.

### Parameter estimation for simulated data under the DLF model

We next evaluate to what extent the two-type DL model can approximate the dynamics of the idealized DLF model, which entails non-independent evolution of gene copies within a family (see methods). We again simulated multiple data sets of 1000 gene families as well as a large data set of 10.000 gene families for the eight-taxon *Drosophila* phylogeny. Note that the total loss rate for a family consisting of one redundant pair will be 2*µ*_*r*_ in the DLF model, whereas the total loss rate for such a family under the two-type DL model (corresponding to the state *X* = (1, 1)) will be *µ*_1_ + *µ*_2_ ≈ *µ*_2_. Because such small families dominate the data, we expect that *µ*_2_ ≈ 2*µ*_*r*_.

In line with this expectation, we find that the estimated value of *µ*_2_ under the two-type DL model corresponds to twice the simulated loss rate per redundant gene (*µ*_*r*_) under the DLF model (fig. S5). Similar observations hold for *v*, although the large posterior variance blurs the expected relationship of *v* in the DLF model to twice *v* in the two-type DL model. Additionally, we note that the posterior mean *µ*_1_ and *λ* values for the two-type DL model are very accurate estimators of the corresponding parameters under the DLF model. For the large data set simulated under the DLF model with *λ* = 0.2, *µ*_*nr*_ = 0.1, *µ*_*r*_ = 3 and *v* = 0.2, we obtain an estimate for the loss rate per excess gene *µ*_2_ of 5.99 (5.35, 6.65), again coinciding with twice the true loss rate per redundant gene (fig. S6). Similarly, the posterior mean estimate for *v* under the two-type DL model is obtained as 0.35 (0.10, 0.62), also approximately corresponding to twice the true value of the underlying DLF model. The much higher posterior variance for this parameter makes the correspondence again however less clear. These simulations therefore suggest it is reasonable to assume that twice the loss rate per *excess* gene, estimated under the two-type DL model, may serve as an approximation to the more intuitive loss rate per *redundant* gene in the DLF model. Posterior predictive simulations for the fitted two-type DL model further show that the posterior predictive distribution is fully compatible with the data simulated under the DLF model (fig. S7), indicating that the two-type DL model indeed can serve as an approximation to the DLF model.

### Analysis of *Drosophila*, yeast and primate data

We performed Bayesian inference for the two-type DL model using gene count data from eight *Drosophila* species, eight yeast species and 11 primates (see methods) and compare the inferred model parameters to parameter estimates for different single-type BDP models (table 1). We note that for the yeast and primates data sets, initial tests indicated that numerical inaccuracies in the likelihood for the two-type model could lead to a failure of the MCMC algorithm to converge in some runs. Increasing the FFT length from *N* = 16 (as employed in our simulations) to *N* = 32 and decreasing tolerance settings in the ODE solver alleviated these issues and resulted in proper convergence. Furthermore, we found that the posterior distribution for the primates data set was bimodal, with one of the modes at *µ*_2_ > 20, *λ* > 1.5 and *r* ≈ 0.2. We deem this unrealistic in light of the results for the single type models and other data sets, signaling the need for a more informative prior. We therefore further restrict *µ*_2_ < 10 for this data set.

**Table 1:** Marginal posterior parameter estimates. For all analyses, the prior on the number of lineages at the root was a Beta-Geometric distribution with *η* and *ζ* parameters fixed to the marginal posterior mean values obtained from the stationary distribution fit (fig. 2, *Drosophila*: *η* = 0.96, *ζ* = 4.01, yeasts: *η* = 0.98, *ζ* = 4.06, primates: *η* = 0.92, *ζ* = 3.23). For the two-type model we use an exponential prior for *µ*_2_ with mean equal to the posterior mean value of *µ* under the single-type model with no extinction. For the primates data we further restrict *µ*_2_ < 10. Other priors are as described in the methods section. *t*_1/2_ and *p*_*x*_ denote the half-life (in My) and probability of x-functionalization (in %) of a type 2 gene respectively

| Parameter | *Drosophila* | Yeasts | Primates (GO:0002376) |
| --- | --- | --- | --- |
| *Single-type (critical)* |  |  |  |
| $\lambda$ | 0.19 (0.19, 0.20) | 0.040 (0.038, 0.041) | 0.18 (0.18, 0.19) |
| *Single-type* |  |  |  |
| $\lambda$ | 0.17 (0.16, 0.17) | 0.022 (0.020, 0.023) | 0.20 (0.19, 0.21) |
| $\mu$ | 0.24 (0.23, 0.25) | 0.059 (0.056, 0.062) | 0.17 (0.16, 0.18) |
| *Single-type (non-extinct)* |  |  |  |
| $\lambda$ | 0.18 (0.17, 0.19) | 0.016 (0.014, 0.019) | 0.24 (0.22, 0.26) |
| $\mu$ | 3.94 (3.71, 4.18) | 1.606 (1.385, 1.813) | 3.46 (3.24, 3.67) |
| *Two-type* |  |  |  |
| $\lambda$ | 0.26 (0.24, 0.28) | 0.027 (0.024, 0.030) | 0.34 (0.32, 0.38) |
| $\mu_1$ | 0.19 (0.18, 0.20) | 0.055 (0.053, 0.058) | 0.13 (0.12, 0.14) |
| $\mu_2$ | 6.37 (5.98, 6.93) | 1.794 (1.638, 1.972) | 4.32 (3.95, 4.71) |
| $\nu$ | 0.20 (0.08, 0.35) | 0.045 (0.007, 0.101) | 0.08 (0.03, 0.14) |
| $r$ | 0.12 (0.10, 0.15) | 0.013 (0.002, 0.031) | 0.04 (0.02, 0.07) |
| $t_{1/2}$ (My) | 11 (10, 12) | 39 (35, 42) | 16 (15, 18) |
| $p_x$ (%) | 3.1 (1.3, 5.0) | 2.3 (0.4, 5.2) | 1.8 (0.7, 3.1) |

Interestingly, the estimated rates for slow processes in the *Drosophila* and primates data sets are roughly on the same scale, corresponding to about 0.1 to 0.4 events per gene per 100 My, whereas the yeast data set yields parameter estimates that are considerably lower. Of course, this is not wholly unexpected, given the vastly different genomic organization of yeast species, with small genomes typically consisting of 5000 to 7000 predicted genes. The overall behavior of parameter estimates across models is similar for all three data sets. We find that the duplication rate estimated for the two-type DL model is higher than for the single-type models, which is in line with our simulations above. The loss rate for an excess (type 2) gene *µ*_2_ is an order of magnitude higher than other rate parameters, with an implied half-life (*t*_1/2_) of a type 2 gene of about 11, 39 and 16 My for the three data sets respectively (table 1). If we assume *µ*_2_ ≈ 2*µ*_*r*_ (see above), these can be interpreted as the half-lives of duplicate *pairs* under the DLF model. Note that the ratios of *λ*/*µ*_2_ of about 0.04, 0.02 and 0.08 are in complete agreement with the Beta-Geometric stationary distribution fit of the single-type model (fig. 2, Appendix 1). The marginal posterior mean estimate of the probability that a new duplicate gets established eventually (i.e. becomes a type 1 gene, rather then suffering loss, *p*_*x*_) is a mere 3% for the *Drosophila* data, about 2% for the yeast data and 2% for the primates data. Again, under the DLF model, this value can be interpreted as the probability that a duplicate *pair* undergoes successful sub- or neofunctionalization so that it is stably established in the genome.

Clearly, a failure to model the different loss rates within a family caused by functional redundancy leads to a loss rate estimate *µ* that is dominated by the large number of non-redundant genes that only rarely get lost. The loss rate in the default single-type BDP model therefore more closely resembles *µ*_1_ than *µ*_2_ in the two-type model. The opposite holds for the single-type model for the number of *excess* genes in non-extinct families. However the loss rates for this model (estimated at 3.94, 1.61 and 3.46 expected loss events per excess gene per 100 My for the three data sets respectively) are considerably lower than the estimates for *µ*_2_ for the two-type model. This is consistent with the dynamics the two-type process is supposed to model, as duplicated genes that sub- or neofunctionalize should pull back the loss rate towards the loss rate of non-redundant genes in the corresponding single-type model. In other words, when assuming all duplicated genes have the same loss rate (as in the single-type non-extinct model), the presence of duplicate genes that have become essential leads to a downwardly biased loss rate when interpreted as the rate of pseudogenization of duplicate genes. An additional explanatory factor may be that the subset of families which do not go extinct has a lower average loss rate per excess gene than the full data set. Lastly, all single-type models seem to underestimate the duplication rate with respect to the two-type model, which may be a result of the biases in the loss rates caused by a failure to separate the dynamics of redundant versus non-redundant duplicated genes.

Posterior predictive simulations indicate that the two-type model provides a better fit to the size distribution than the single-type model for all data sets (fig. 4, fig. S8, fig. S10, fig. S11). For all three data sets we find that the Kullback-Leibler (KL) divergence 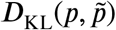 from the posterior predictive frequency distribution 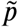 to the observed frequency distribution *p* for the two-type model is less than half of the same KL divergence obtained under the default single-type model (fig. S8, fig. S9). We note that the different degrees of correspondence of the posterior predictive distributions to the observed size frequency distributions for different taxa suggests rate heterogeneity across *branches* of the species tree, a complication we ignore in the present paper. As expected, the posterior predictive size distribution is underdispersed with respect to the true distribution further in the tail (fig. S12), which is a consequence of ignoring rate heterogeneity across *families*. This is less so for the single-type model, where the tail of the posterior predictive distribution is closer to the observed distribution. This is simply a result of the distribution on the number of lineages at the root being better preserved under the process, which has a lower overall event rate. We note that the posterior predictive distribution for the non-zero counts under the two-type model is nearly indistinguishable from the posterior predictive distribution for the single-type model for non-extinct families.

**Figure 4:**
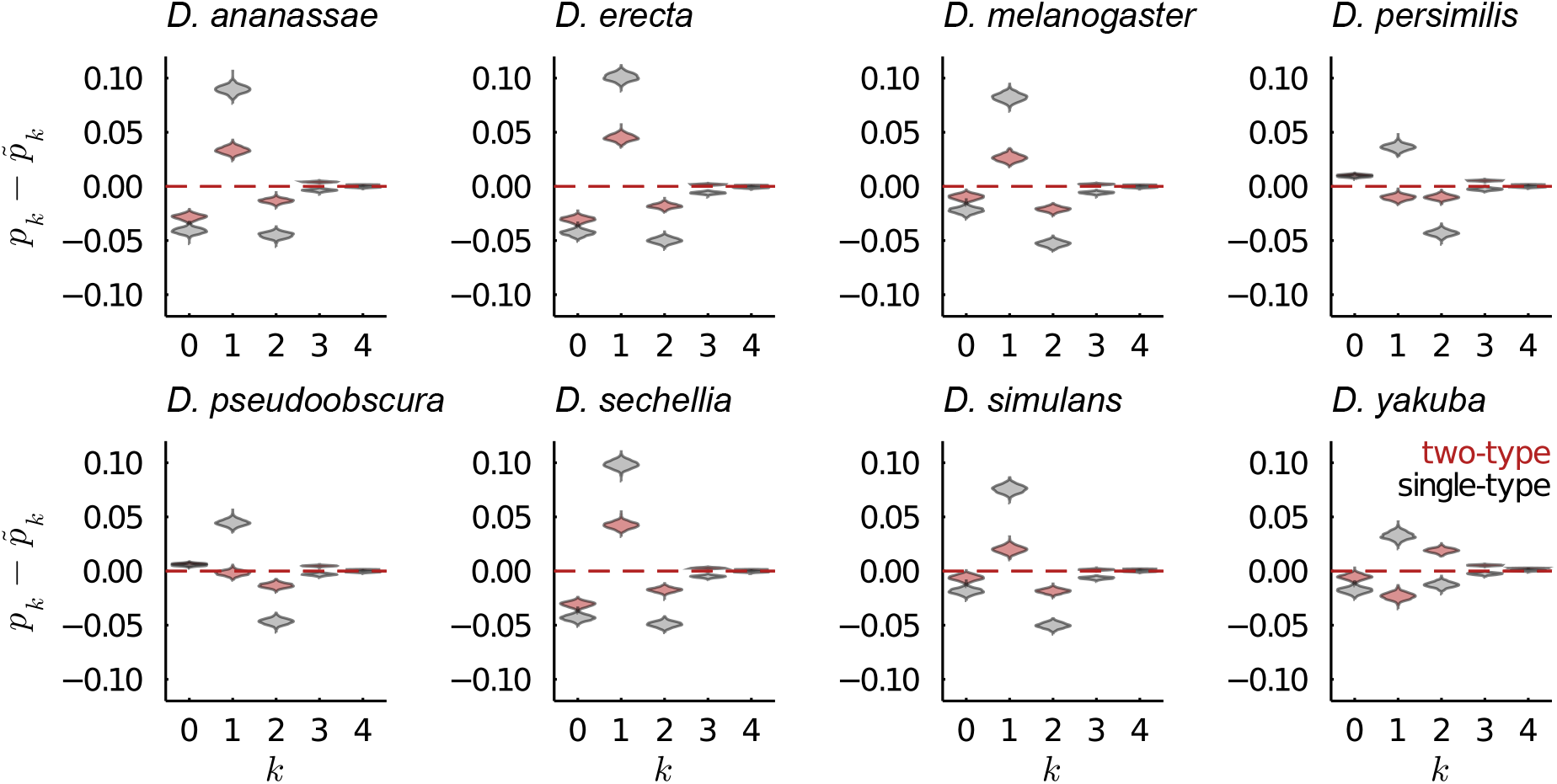
Posterior predictive densities of the gene family size distributions for each of the eight leaves of the *Drosophila* phylogeny. The distribution of the differences of the observed gene family size frequencies (*p*_*k*_) with the frequencies observed in 1000 posterior predictive simulations 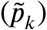 is plotted. In red and gray posterior predictive distributions for the two-type and single-type model are shown respectively.

While rate heterogeneity across families could be accounted for by using a mixture modeling approach, we refrain from doing so for reasons of computational feasibility. We may however apply an *ad hoc* procedure to apply rate heterogeneity across families in our posterior predictive simulations by considering the fit of the BG distribution to the data. As we discuss in the methods section and Appendix 2, the BG distribution is the stationary distribution of a special case of the two-type DL model where *µ*_1_ = *v* = 0 and where the ratio *λ*/*µ*_2_ is iid distributed according to a Beta distribution across families. The parameters of this stationary distribution can be easily estimated from the data, and we estimated the dispersion parameter for the *Drosophila* data for instance at *ζ* = 4.01 (fig. 2). If we assume *λ*/*µ*_2_ to be Beta distributed across families, and take for each replicate simulation of the posterior distribution the value of *λ*/*µ*_2_ as the mean of this Beta distribution, we may simulate from the posterior distribution under this assumption as follows: For each of the *i* ∈ (1, …, 1000) replicate simulations we sample a (*λ*/*µ*_2_, *µ*_2_)_*i*_ pair from the posterior and compute *α*_*i*_ = *ζ* (*λ*_*i*_/*µ*_2,*i*_) and *β*_*i*_ = *ζ* (1−*λ*_*i*_/*µ*_2,*i*_). For each of the *j* ∈ (1, …, *n*) families in the *i*th simulation replicate, we sample a random value for*ξ*_*ij*_ = *λ*_*ij*_/*µ*_2,*ij*_ by sampling from Beta(*α*_*i*_, *β*_*i*_). We then obtain *λ*_*ij*_ and *µ*_2,*ij*_ by assuming *µ*_2,*ij*_ = *µ*_2,*i*_ and *λ*_*ij*_ = *ξ*_*ij*_*µ*_2,*i*_. Note that this procedure translates the assumed variation in *λ*/*µ*_2_ across families to heterogeneity in *λ* alone across families. The joint posterior distribution of the family-specific rates obtained following this procedure is shown in fig. 5. The resulting posterior predictive distribution still fits the small family sizes well (as in fig. 4), but now no longer underestimates the proportions of larger families, and yields predictions compatible with the observed power law tail (fig. 5). This again clearly shows that the power-law tail of the gene family size distribution can be explained by rate heterogeneity across families.

## Discussion

We describe a two-type continuous-time branching process model of gene family evolution, show the feasibility of estimating its parameters from incomplete data in a phylogenetic context using simulations, and perform Bayesian inference of model parameters for comparative genomic data sets. Comparisons with closely related single-type phylogenetic BDP models highlight the shortcomings of these models and indicate how a two-type process may provide a first step towards more realistic stochastic models of gene family evolution, providing more detailed and biologically meaningful quantitative insights in the associated evolutionary processes.

The main virtue of the two-type DL model is the discrimination between non-redundant functional genes that are only rarely lost, and duplicates which may be largely redundant and therefore more prone to loss by pseudogenization. We noted that the model proposed here can be viewed as an approximation to the gene family evolution dynamics where duplicated genes are (partially) functionally redundant and evolve in a non-independent manner (as in the DLF model). Compared to the DLF model, the loss rate *µ*_2_ in the two-type process does *not* correspond to the loss rate per redundant gene, but rather the loss rate per *excess* gene in a family. We explicitly examined the connection between the two models using simulated data and find that parameter estimates under the two-type DL model can be interpreted in the context of the more intuitive DLF model.

An alternative approach to account for differences in loss rates within a family which may be natural to adopt is to consider an age-dependent process (more precisely a ‘budding’ age-dependent branching process (Greenman and Chou 2016)), where the loss rate of a duplicated gene decreases through time. Such a model has been considered by Zhao et al. (2015), albeit not in the context of gene content evolution along a phylogeny. Inference of model parameters relies however on knowledge of the gene family trees and ages of duplication events (dated reconciled gene trees in the comparative genomic setting), the estimation of which is of course an extremely challenging statistical problem in its own right. Without access to such high-quality gene trees, inference from gene counts alone seems highly challenging. Indeed, statistical inference of age-dependent branching processes more generally is an active research topic of considerable mathematical sophistication (e.g. Fok and Chou 2013; Greenman and Chou 2016).

It is important to stress what is *not* modelled by the phylogenetic two-type DL process. Most importantly, we do not explicitly model the population genetics of fixation of copy number variants (CNVs). Clearly, most duplication events will either be lost due to drift if neutral, or lost by purifying selection when deleterious, and as a result leave no trace in extant genomes. The duplication rate *λ* in the model should therefore be interpreted as a proxy for the rate at which gene duplications that rise to high frequency occur. We have however no guarantee that the observed copy numbers are monomorphic in their respective populations, so the presence of low-frequency CNVs in the data may cause *λ* to overestimate the rate at which duplications that fix in the population occur. Similar considerations hold for the loss rate parameters *µ*_1_ and *µ*_2_, which should be interpreted as rates of gene deletion events, not the rates of loss of unfixed duplicate genes by genetic drift or purifying selection. Of course, exactly the same issues hold for single-type models as well. We note that, if gene duplications were predominantly neutral (which is of course extremely unlikely), the estimated *λ* should roughly correspond to the per-gene duplicative mutation rate. For *Drosophila*, the duplicative mutation rate has been estimated at 1.25 × 10^−7^ duplications per gene per generation (Schrider et al. 2013).

Taking this as a crude estimate of the order of magnitude of the duplicative mutation rate, and considering a long-term average generation time between 7 to 20 days, the expected duplication rate *λ* under neutrality would be *at least* a 100 times larger than the rates we estimated under the two-type DL model, suggesting that the vast majority of duplications is deleterious. *λ* more detailed picture of the distribution of fitness effects of new duplications remains however elusive.

The two-type DL model provides new quantitative insights in the long-term evolutionary dynamics of duplicated genes. In our analyses of comparative genomic data sets, we find that the loss rate per excess gene is much higher than the loss rate per base gene, suggesting that even when gene duplicates establish in a population, the selective pressures ensuring their maintenance differ strongly from those maintaining typical single-copy genes. For instance, for the *Drosophila* genus, we estimated that on average half of the duplicated genes are maintained over a period of approximately 11 My and that about 3% of the duplicated genes may eventually become stably established (complete x-functionalization) so that their loss rates reflect those of base genes. The half-life of a typical base gene on the other hand is estimated at 371 My. Concomitantly, our results indicate that estimates of long-term gene loss rates based on simple BDP models (as in e.g. Hahn, Han, and Han (2007) for *Drosophila*) likely underestimate the actual loss rates of duplicated genes significantly, while estimates of duplication rates tend to be in rough agreement with those presented here.

What then causes gene duplicates that establish in a population to remain prone to higher loss rates? Or conversely, how are gene duplicates that are non-essential and prone to loss at relatively high rates established in the first place? Of course, completely redundant duplicates may drift to fixation, and it is unsurprising that such genes would be prone to higher loss rates than their essential counterparts (Walsh 2003). Many gene duplicates that rise to fixation may however be adaptive (Han et al. 2009; Innan and Kondrashov 2010; Kondrashov 2012), demanding an answer to the first question. As Kondrashov (2012) stressed, it may be misleading to think of gene duplicates as either completely redundant or non-redundant, as many adaptive duplications may establish as a consequence of positive selection for increased dosage in a stressful environment, despite being *qualitatively* redundant. In such cases, a changing environment or the emergence of other genetic variants may alter selection pressures over time, and while some duplicated gene may promote adaptation during some environmental challenge, it may return to a state of complete redundancy or even come at a fitness cost later (Kondrashov 2012). Our phylogenetic analyses may go some way supporting this view of long-term gene family evolution, where many of the duplicated genes in multi-copy gene families reside for some time in the genome, but eventually suffer loss before undergoing complete sub- or neofunctionalization.

## Acknowledgements & Funding

Arthur Zwaenepoel acknowledges funding from the flemish fund for scientific research (FWO). Yves Van de Peer acknowledges funding from the European Research Council (ERC) under the European Union’s Horizon 2020 research and innovation program (grant agreement No 833522) and from Ghent University (Methusalem funding, BOF.MET.2021.0005.01).

## Supplementary material

## Appendix 1: Transition probabilities for the two-type DL model

Here we derive the probability generating function (pgf) for the two-type DL process defined in the main text using techniques from Bailey (1990) and Xu et al. (2015). Let *f*_*ij*_(*s*_1_, *s*_2_, *t*) denote the pgf for the two-type branching process model in eq. 1 of the main text when starting at *X*(0) = (*i, j*), i.e.

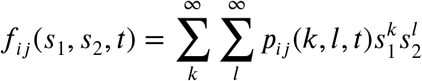

Importantly, the branching property implies the following relationship among the pgfs for different *X*(0)

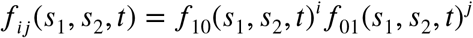

(e.g. Athreya and Ney (1972)), so we may work with *f*_10_ and *f*_01_ and recover the desired pgfs easily. Let *p*_*ij*_(*k, l, dt*) = *r*_*ij*_(*k, l*)*dt* + *o*(*dt*), we also define the auxilliary generating functions

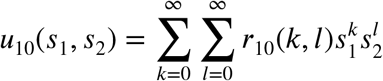

and *u*_01_(*s*_1_, *s*_2_) analogously in terms of *r*_01_(*k, l*). Note that the *r*_10_ and *r*_01_ functions are determined by the definition of the process in terms of its infinitesimal rates in eq. 1. Filling in the relevant parameters we obtain

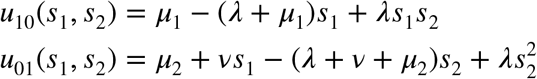

The pgfs for the process are related to the auxilliary generating functions

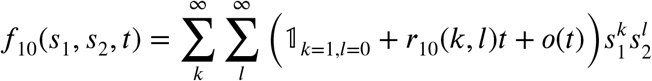

and similarly for *f*_01_, so that we get

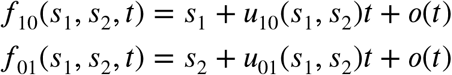

Note furthermore that *∂f*_*ij*_(*s*_1_, *s*_2_, *t*)/ *∂t* = *u*_*ij*_(*s*_1_, *s*_2_). We can expand *f*_10_ as a Taylor series around *t*

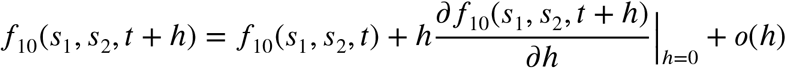

Now exploiting the property of the branching process that

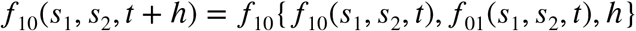

we can rewrite the Taylor expansion as

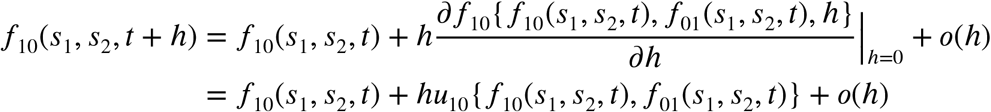

Which shows that

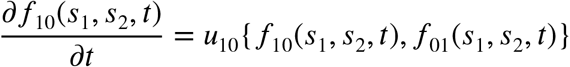

and analogously for *∂f*_01_/*∂t*. Combining this result with the generating functions *u*_10_ and *u*_01_, we arrive at the following system of non-linear ordinary differential equations (ODEs)

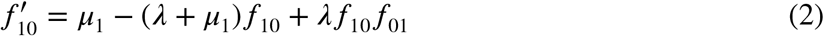

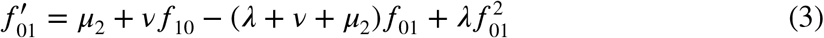

Where the arguments *s*_1_, *s*_2_ and *t* are omitted for notational convenience and differentiation is with respect to *t*.

## Appendix 2: The stationary distribution of a birth-death process with no extinction and rate heterogeneity across families

We propose the Beta-geometric distribution as a reasonable prior for the number of ancestral lineages. This choice is motivated by two main factors: (1) the observed size distribution of gene families shows an approximate power-law decay and (2) the stationary distribution of a special case of the two-type process with duplication and loss rate heterogeneity across families can be shown to lead to a Beta-geometric probability mass function (pmf).

Consider the two-type branching process model proposed in the main text, where we set *µ*_1_ = *v* = 0 and *µ*_2_ = *µ*, and where we assume the process starts with a single type 1 gene. In this model, the number of type 1 genes stays 1 perpetually, while the number of type 2 genes evolves as a stochastic birth-death-immigration process (BDIP) with immigration rate *λ*. The system of differential-difference equations for the transient distribution of the number of type 2 genes is

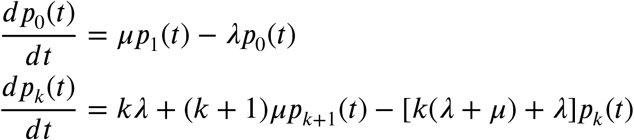

Setting *dp*_*k*_(*t*)/*dt* = 0, the stationary pmf *π* can be obtained as

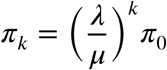

With the constraint 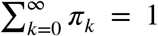 and under the assumption that *λ < µ*, we can find that *π*_0_ = 1 − *λ*/*µ* so that the stationary pmf under this model becomes

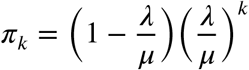

Being a Geometric distribution on *k* ≥ 0 with parameter 1 − *λ*/*µ*. Note that the distribution for the *total* number of genes is this Geometric distribution shifted by one, *π*_*k*_ = (1 − *λ*/*µ*)(*λ*/*µ*)^*k*−1^, *k >* 0.

Now (considering point (1) above), empirical size distributions of non-extinct gene families are overdispersed with respect to the Geometric stationary prediction under this model. If however duplication and loss rates vary across families (which is of course expected to be the rule rather than the exception given the many functional and structural features of genes that should affect their propensity to duplicate or pseudogenize), the model above with Geometric stationary distribution gives rise to power-law like distributions for some models of rate heterogeneity.

Consider now *λ* and *µ* for each gene family to be a random variable iid from some distribution. A particularly straightforward and flexible model is to assume 1 − *λ*/*µ* to be distributed according to a Beta(*α, β*) distribution across families (or equivalently, *λ*/*µ* ∼ Beta(*β, α*)). The resulting compound stationary distribution becomes a so-called Beta-Geometric BG(*α, β*) distribution with pmf

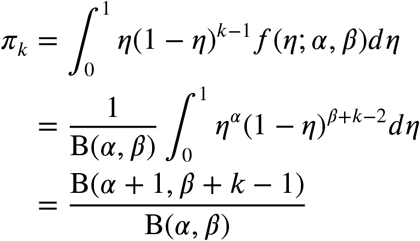

for *k >* 0 and where *f* (*η*; *α, β*) = B(*α, β*)^−1^*η* ^*α*−1^(1 − *η*)^*β* −1^ is the density of the Beta distribution. The mean of this distribution is (*α* + *β*)/ *α*, and the expected value of *λ*/*µ* across families under this model is (*α* + *β*)/*β*. In our work, we mostly employ the alternative parameterization using the mean *η* = *α*/(*α* + *β*) and dispersion parameters *ζ* = *α* + *β* of the underlying Beta distribution.

Having a meaningful interpretation as the stationary distribution of a simple birth-death process with rate heterogeneity, the BG distribution provides a flexible two-parameter alternative to the Geometric distribution employed by Rabier, Ta, and Ané (2014), Zwaenepoel and Van de Peer (2019) and Zwaenepoel and Van de Peer (2020) which provides an excellent fit to observed gene family size distributions. In addition to serving as an apt prior for the number of ancestral genes in a family, a fit of the BG distribution to observed data provides information on the ratio of *λ*/*µ* (*λ*/*µ*_2_ in the context of the two-type process) and the variation of this ratio across families. We infer parameters of the BG distribution using a uniform prior on *η* and flat improper prior on *ζ*, using Turing.jl (Ge, Xu, and Ghahramani 2018) with the NUTS sampler (Hoffman and Gelman 2014).

## Appendix 3: Inference for the single-type model with the Beta-Geometric prior distribution

For the single-type linear birth-death process model we use the same Beta-Geometric prior distribution for the number of genes at the root as for the two-type process. We implemented this in the `DeadBird` package for statistical inference of linear phylogenetic birth-death-immigration processes (which builds on work in Zwaenepoel and Van de Peer (2020)), which uses the algorithm of Csűrös and Miklós (2009) to compute the data log-likelihood. The algorithm of Csűrös and Miklós (2009) makes use of *conditional survival likelihoods*, and is exact, in the sense that it does not require a truncation of the state space. The conditional survival likelihood is defined as

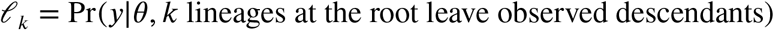

Note that the maximum *k* is bounded by the observed data (Csűrös and Miklós 2009), so that 1 ≤ *k* ≤ *b* where *b* is the relevant bound. Csűrös and Miklós (2009) describe an efficient algorithm to compute *l_k_*. Given a prior on the number of extant genes at the root *Z*^(*a*)^, the complete data likelihood can be obtained as

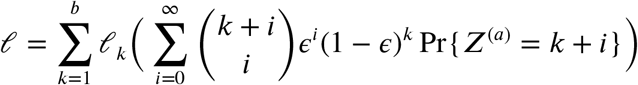

Where *ϵ* is the extinction probability at the root, i.e. the probability that a lineage extant at the root leaves no observed descendants.

For a Geometric(*η*) prior Pr{*Z* = *k*} = *η* (1 − *η*)^*k*−1^, it is easy to show that

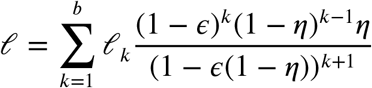

For the Beta-Geometric prior distribution we did not obtain a closed form solution. Using the property of the Beta function that 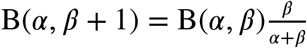 we can find

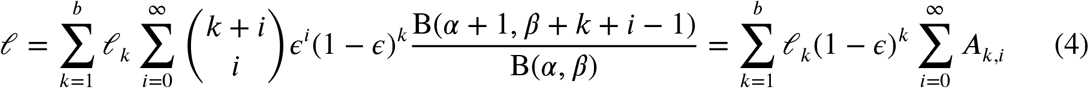

With the recusion relations

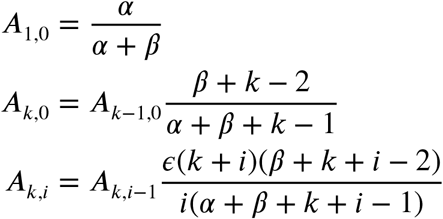

Which allows to efficiently approximate the infinite series in (4) above by some partial sum sufficiently far in the series. The sum converges very rapidly, so that in practice no more than 10 terms are needed. Similar formulas hold in the case where the domain of the Beta-Geometric distribution is *k* ≥ 0. Note that the sum in (4) reduces to the result for the Geometric distribution with *η* = *α* /(*α* + *β*) as (*α* + *β*) →∞.

To compute the condition factor required for correcting the likelihood in accord with the data filtering strategy, we require the probability of a gene family going extinct given the extinction probability of a single lineage *ϵ*. This probability can be obtained as

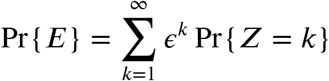

Which can be easily shown to correspond to *ηϵ*/(1 − *ϵ*(1 − *η*)) for the Geometric distribution on *Z*^(*a*)^. For the Beta-Geometric distribution, a little more work leads to

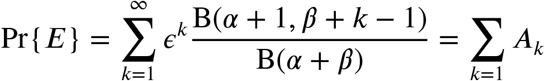

with the recursion relation

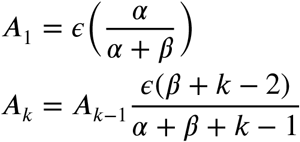

So that we can efficiently approximate the infinite sum by some partial sum sufficiently far in the series (*k >* 100, say). Together these results allow the application of a Beta-Geometric prior distribution on the number of lineages at the root in the context of the algorithm of Csűrös and Miklós (2009).

## Supplementary figures & tables

**Figure S1:**
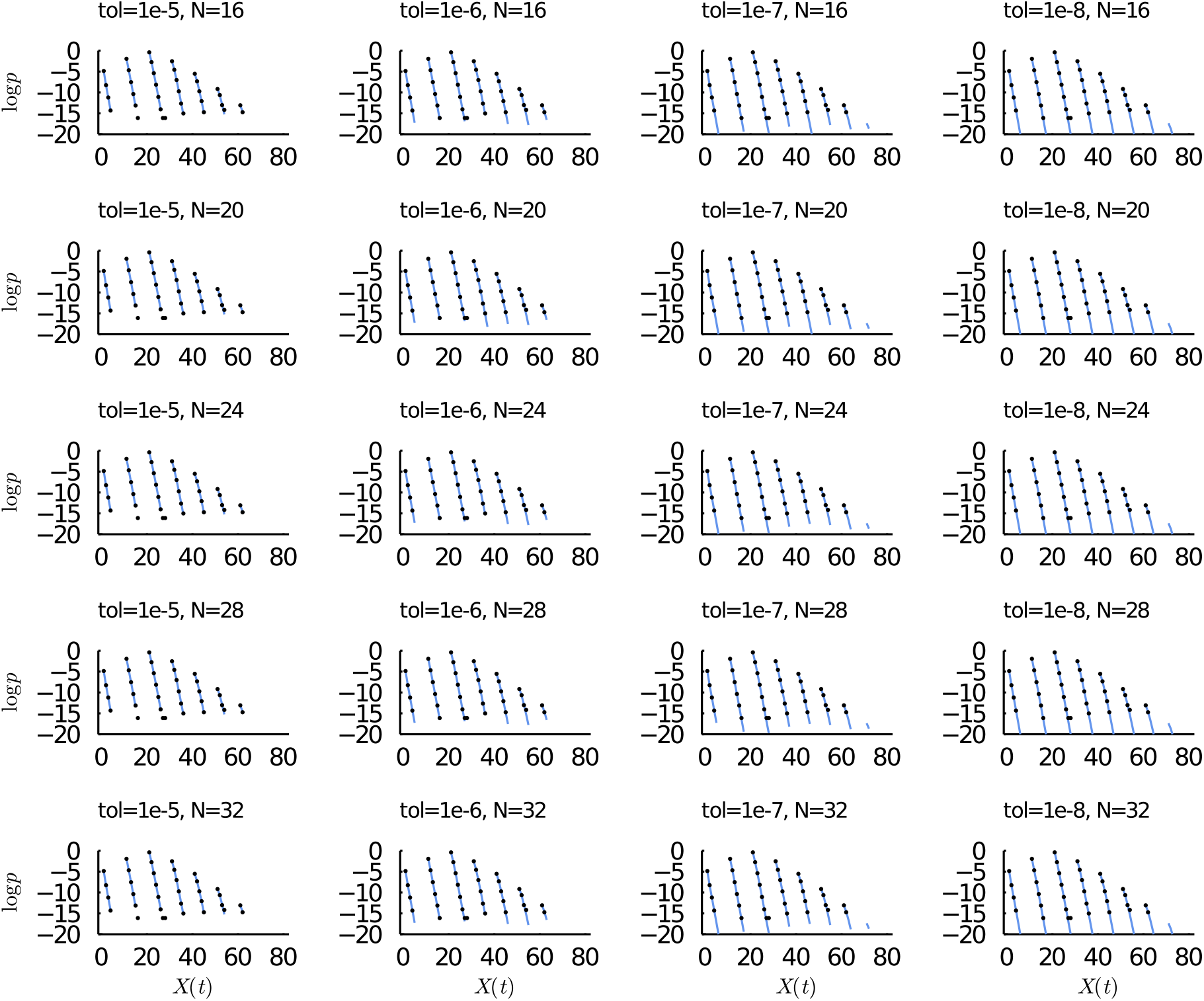
Comparison of log transition probabilities computed using the pgf method (blue lines) and estimated using Monte Carlo simulations (black dots) for different algorithmic settings (tolerance settings in the ODE solver (tO1) and FFT length *N*). Parameters were *λ* = 0.2, *µ*_1_ = 0.1, *ν* = 0.2 and *µ*_2_ = 5. Transition probabilities are computed from the state *X*(0) = (2, 3) to states *X*(*t*) = (*i, j*) where 0 ≤ *i <* 8, 0 ≤ *j <* 10 and *t* = 1. Target states along the *x*-axes are ordered [(0, 0), (0, 1), …, (0, 9), (1, 0), (1, 1) …]. Monte Carlo estimates are based on 10 million independent simulations from the model.

**Table S1:** Taxon names, data sources and ‘accession’ strings for sequence data used in this study.

| id | taxon | source | accession |
| --- | --- | --- | --- |
| dere | *Drosophila erecta* | NCBI | GCF_003286155.1_DereRS2 |
| dana | *Drosophila ananassae* | NCBI | GCF_003285975.2_DanaRS2.1 |
| dsim | *Drosophila simulans* | NCBI | GCF_000754195.2_ASM75419v2 |
| dper | *Drosophila persimilis* | NCBI | GCF_003286085.1_DperRS2 |
| dpse | *Drosophila pseudoobscura* | NCBI | GCF_009870125.1_UCI_Dpse_MV25 |
| dmel | *Drosophila melanogaster* | NCBI | GCF_000001215.4_Release_6_plus_ISO1_MT |
| dyak | *Drosophila yakuba* | NCBI | GCF_000005975.2_dyak_caf1 |
| dsec | *Drosophila sechellia* | NCBI | GCF_004382195.1_ASM438219v1 |
| Ecymbalariae | *Eremothecium cymbalariae* | YGOB, v7-Aug2012 | Ecymbalariae |
| Egossypii | *Eremothecium gossypii* | YGOB, v7-Aug2012 | Egossypii |
| Klactis | *Kluyveromyces lactis* | YGOB, v7-Aug2012 | Klactis |
| Lkluyveri | *Lachancea kluyveri* | YGOB, v7-Aug2012 | Lkluyveri |
| Lthermotolerans | *Lachancea thermotolerans* | YGOB, v7-Aug2012 | Lthermotolerans |
| Lwaltii | *Lachancea waltii* | YGOB, v7-Aug2012 | Lwaltii |
| Tdelbrueckii | *Torulaspora delbrueckii* | YGOB, v7-Aug2012 | Tdelbrueckii |
| Zrouxii | *Zygosaccharomyces rouxii* | YGOB, v7-Aug2012 | Zrouxii |
| Hsapi | *Homo sapiens* | Ensembl, release 102 | Homo_sapiens.GRCh38 |
| Ptrog | *Pan troglodytes* | Ensembl, release 102 | Pan_troglodytes.Pan_tro_3.0 |
| Pabel | *Pongo abelii* | Ensembl, release 102 | Pongo_abelii.PPYG2 |
| Nleuc | *Nomascus leucogenys* | Ensembl, release 102 | Nomascus_leucogenys.Nleu_3.0 |
| Mmula | *Macaca mulatta* | Ensembl, release 102 | Macaca_mulatta.Mmul_10 |
| Panub | *Papio anubis* | Ensembl, release 102 | Papio_anubis.Panu_3.0 |
| Csaba | *Chlorocebus sabaeus* | Ensembl, release 102 | Chlorocebus_sabaeus.ChlSab1.1 |
| Cjacc | *Callithrix jacchus* | Ensembl, release 102 | Callithrix_jacchus.ASM275486v1 |
| Csyri | *Carlito syrichta* | Ensembl, release 102 | Carlito_syrichta.Tarsius_syrichta-2.0.1 |
| Mmuri | *Microcebus murinus* | Ensembl, release 102 | Microcebus_murinus.Mmur_3.0 |
| Pcoqu | *Propithecus coquereli* | Ensembl, release 102 | Propithecus_coquereli.Pcoq_1.0 |
| Ogarn | *Otolemur garnettii* | Ensembl, release 102 | Otolemur_garnettii.OtoGar3 |

**Figure S2:**
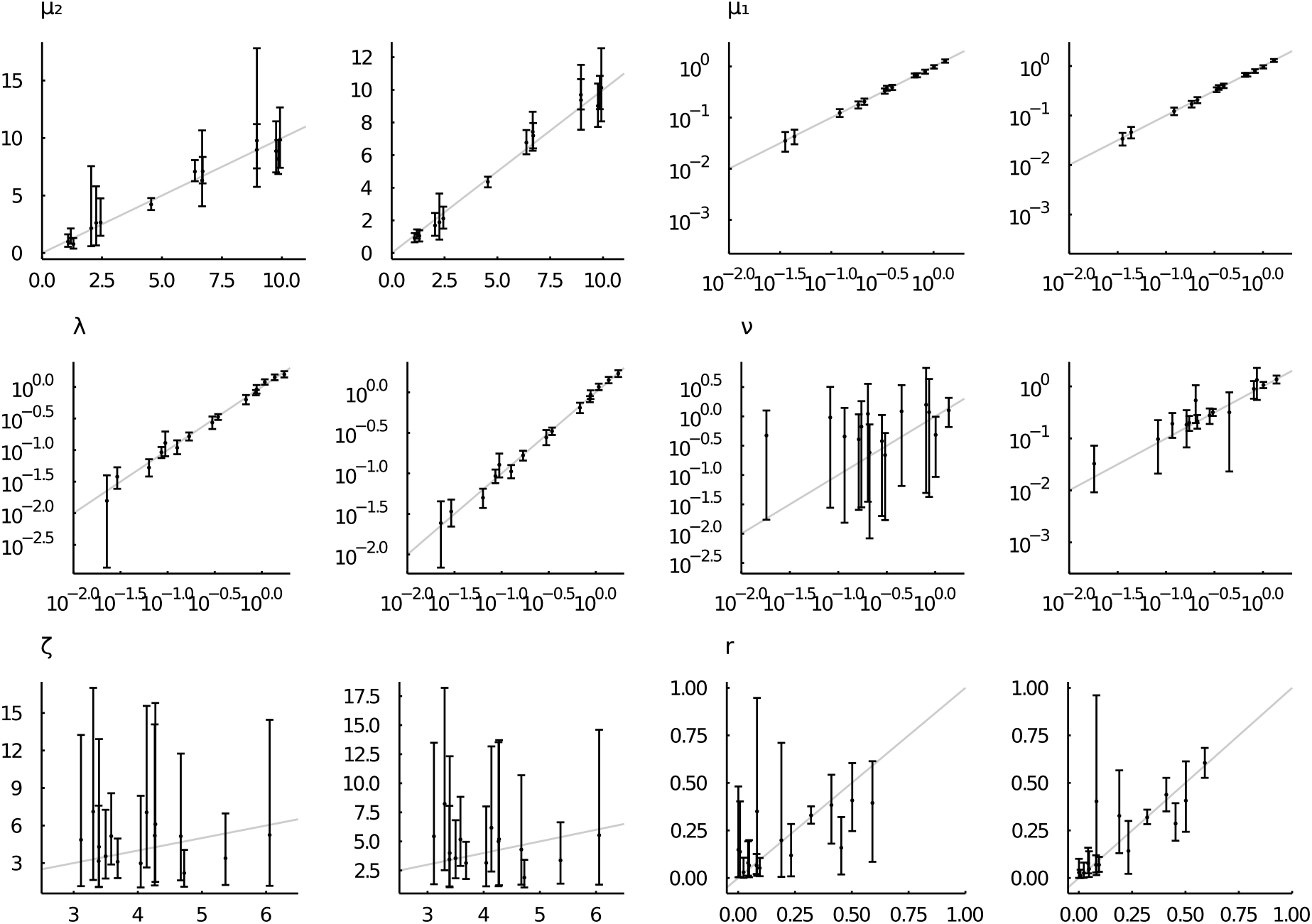
Posterior mean parameter estimates and 95% uncertainty intervals (*y*-axis) as a function of the true simulated value (*x*-axis) for simulations of data sets of 1000 gene families. *µ*_2_ values were drawn uniformly from the interval (1, 10), while *λ*/*µ*_2_ *µ*_1_/*µ*_2_ and *v*/*µ*_2_ were drawn from a Beta(9, 1) distribution. *r* was drawn from a Beta(3, 1) distribution and *ζ* from a log normal distribution with mean 3 and variance 0.2. Plots are shown in pairs, with on the left the results conditioning on the incompletely observed data *Z* = *X*_1_ + *X*_2_ while on the right results conditioning on the fully observed data *X* = (*X*_1_, *X*_2_) are shown.

**Figure S3:**
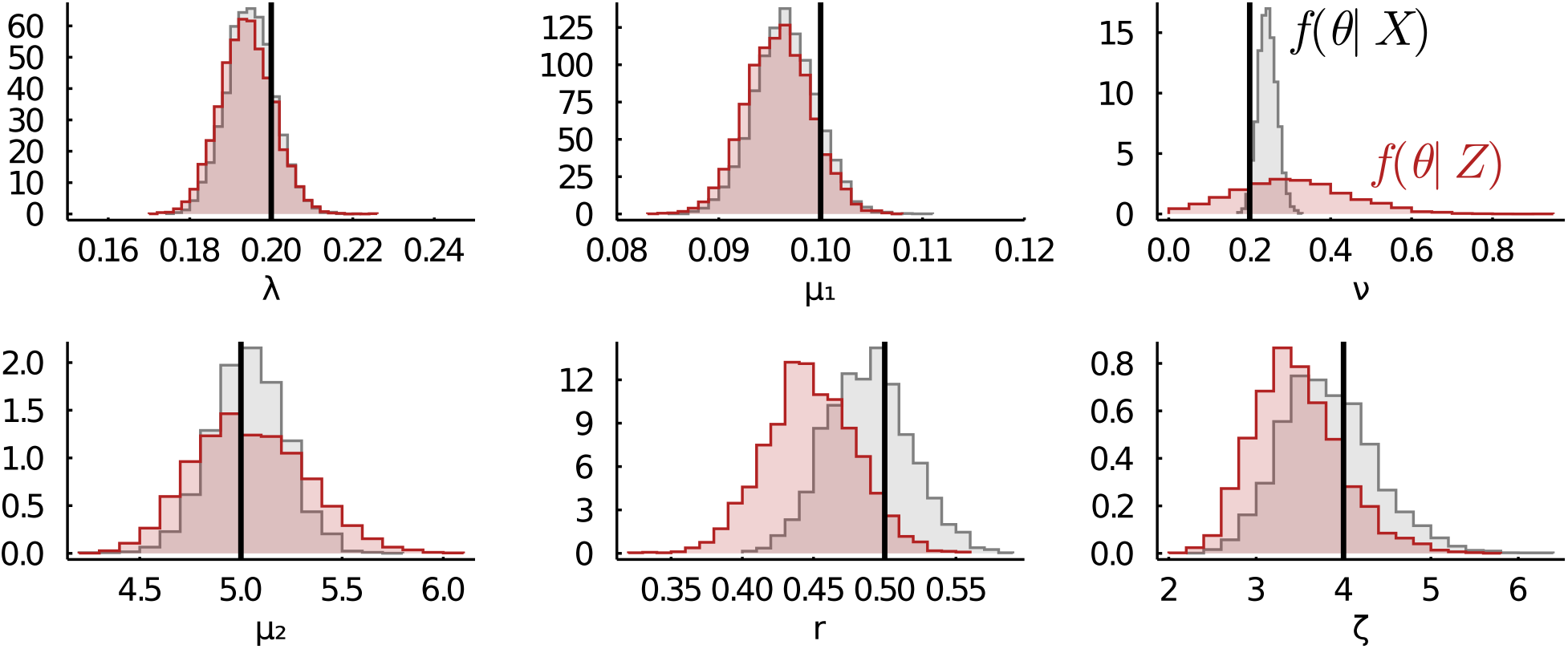
An independent replicate of the simulation shown in fig. 3

**Figure S4:**
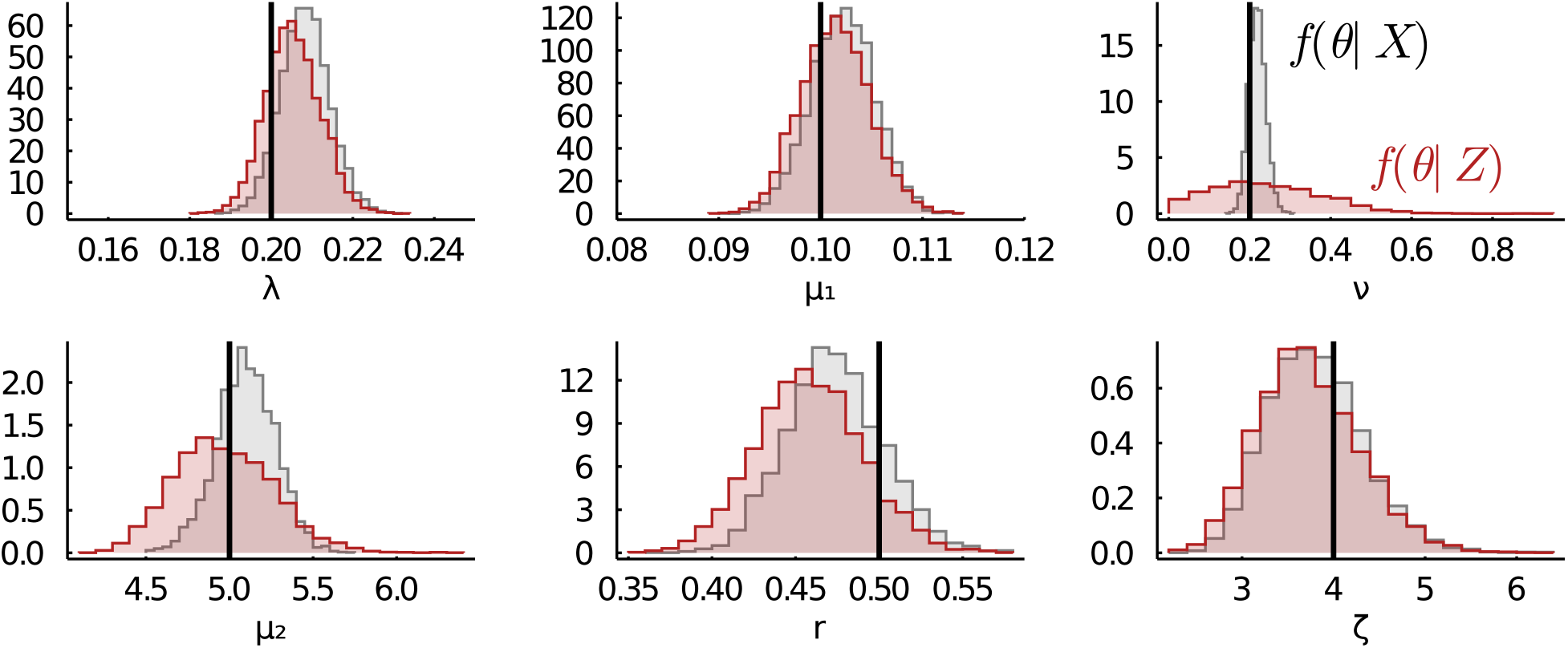
An independent replicate of the simulation shown in fig. 3

**Table S2:** Posterior distribution summary showing marginal posterior means and 95% uncertainty intervals (lower, upper) for various models for the simulation replicate associated with fig. 3.

| model | parameter | mean | lower | upper |
| --- | --- | --- | --- | --- |
| Single-type | $\lambda$ | 0.14 | 0.14 | 0.15 |
| | $\mu$ | 0.12 | 0.11 | 0.13 |
| | $\zeta$ | 6.77 | 4.62 | 9.96 |
| Single-type (fixed $\zeta$) | $\lambda$ | 0.14 | 0.14 | 0.15 |
| | $\mu$ | 0.12 | 0.11 | 0.12 |
| Single-type (non extinct) | $\lambda$ | 0.14 | 0.13 | 0.15 |
| | $\mu$ | 1.25 | 1.11 | 1.42 |
| | $\zeta$ | 3.69 | 2.65 | 5.11 |
| Single-type (non extinct fixed $\zeta$) | $\lambda$ | 0.14 | 0.14 | 0.15 |
| | $\mu$ | 1.27 | 1.13 | 1.41 |
| Two-type | $\lambda$ | 0.19 | 0.18 | 0.21 |
| | $\mu_1$ | 0.10 | 0.09 | 0.11 |
| | $\nu$ | 0.25 | 0.03 | 0.54 |
| | $\mu_2$ | 4.94 | 4.44 | 5.51 |
| | $\zeta$ | 4.20 | 3.15 | 5.60 |
| | $r$ | 0.48 | 0.41 | 0.54 |

**Figure S5:**
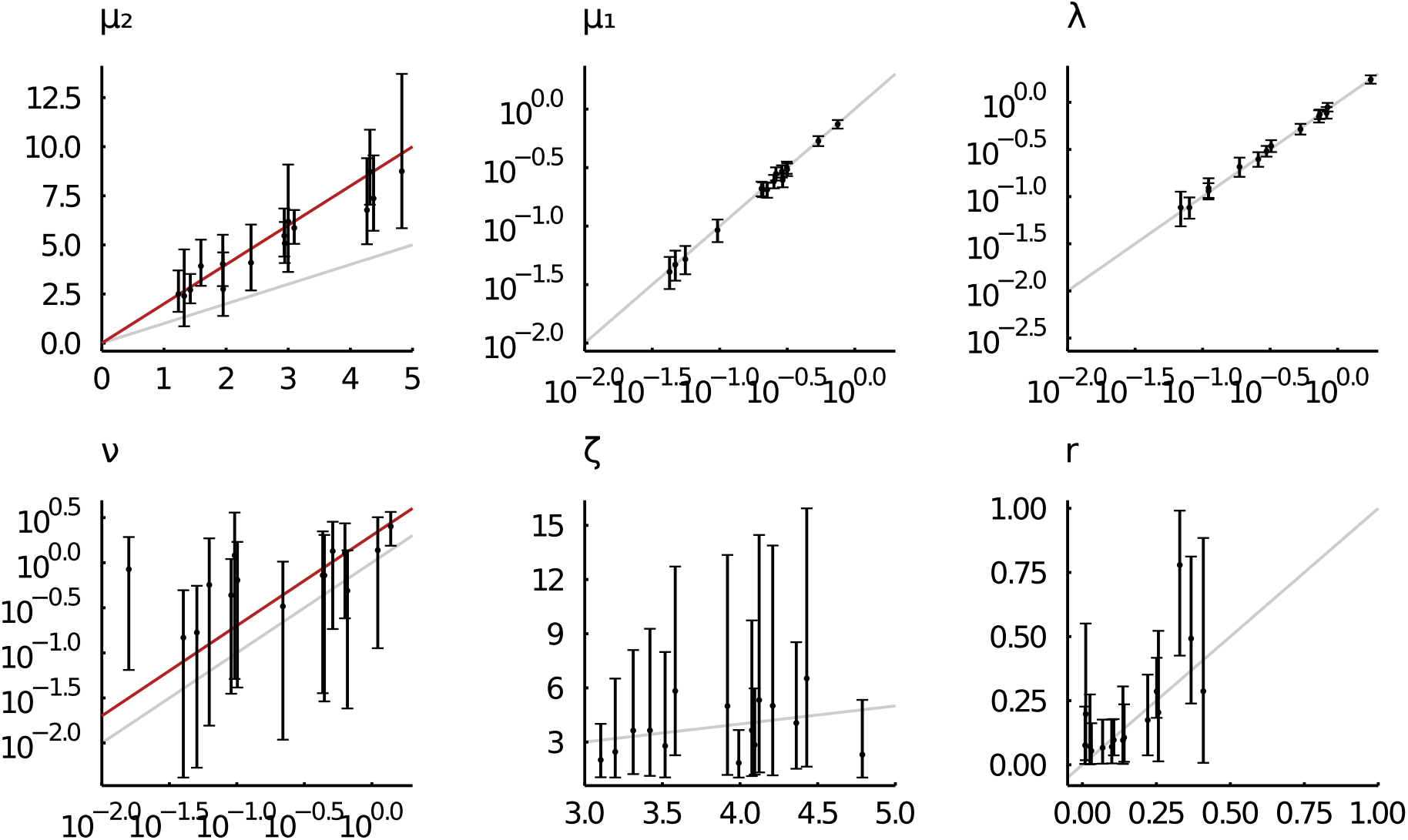
Posterior mean parameter estimates and 95% uncertainty intervals (*y*-axis) as a function of the true simulated value (*x*-axis) for simulations of data sets of 1000 gene families. Data sets were simulated under the DLF model, while inference was performed under the two-type DL model. *µ*_2_ values were drawn uniformly from the interval (1, 5), while *ζ*/*µ*_2_ *µ*_1_/*µ*_2_ and *ν* /*µ*_2_ were drawn from a Beta(9, 1) distribution. *r* was drawn from a Beta(3, 1) distribution and *ζ* from a log normal distribution with mean 3 and variance 0.2. Inference is based on the total gene count (*Z*). The red lines for *µ*_2_ and *v* mark the expected approximate relationship between the simulated parameter value in the DLF model and the corresponding parameter in the two-type DL model (i.e. *µ*_2_ ≈ 2*µ*_*r*_ and *ν* ≈ 2 *ν* _DLF_).

**Figure S6:**
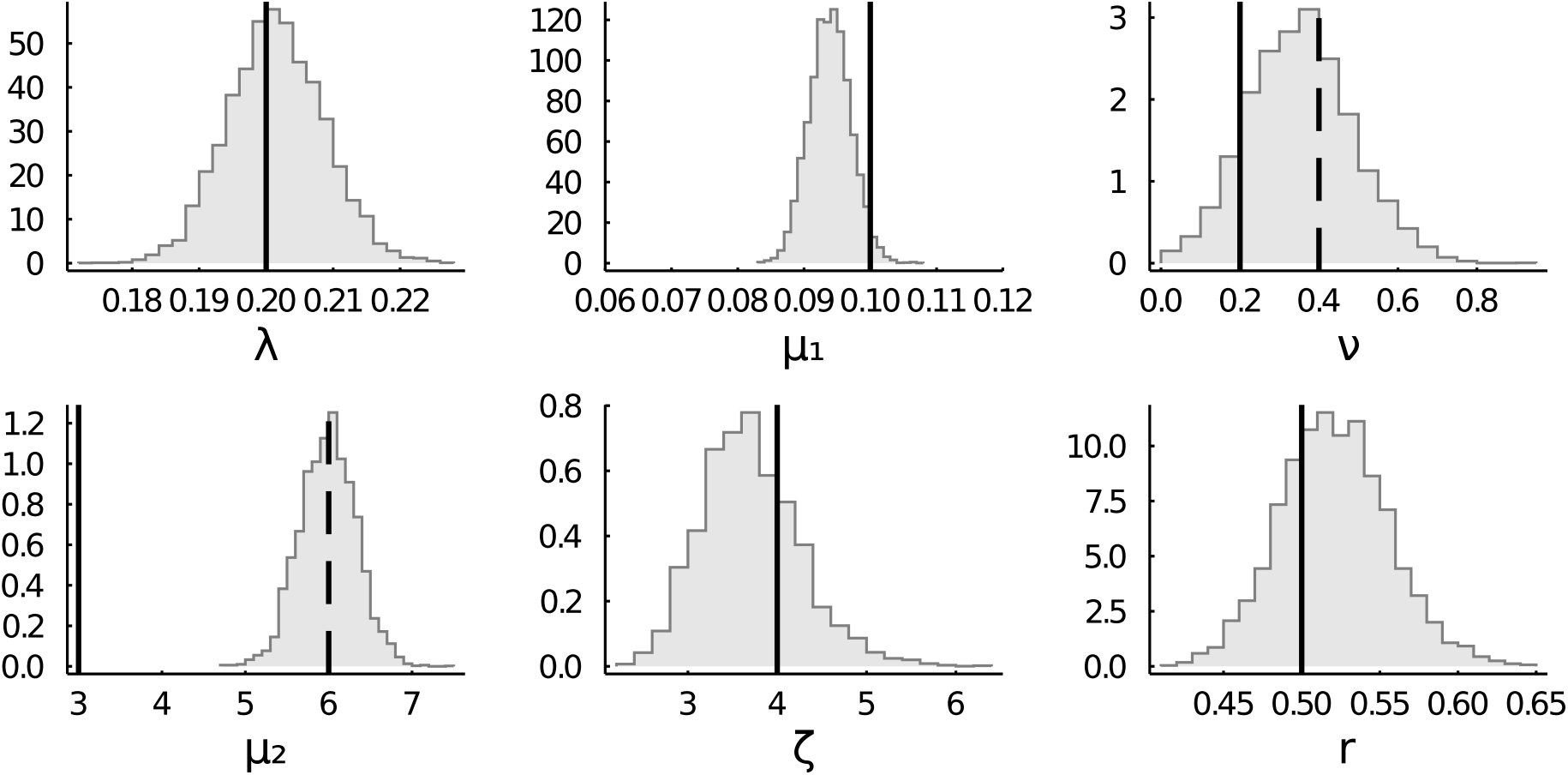
Marginal posterior distributions for the two-type DL model applied to data simulated under the DLF model. Vertical solid lines mark the true simulated values, while dashed lines mark the expected approximate correspondences *µ*_2_ ≈ 2*µ*_*r*_ and *ν* ≈ 2 *ν* _DLF_ (see main text).

**Table S3:** Summary of the posterior distributions showing marginal posterior means and 95% uncertainty intervals (lower, upper) for various single-type models fitted to the data simulated from the DLF model (fig. S6). S8, but now for each species separately. Results for the two-type model are displayed in red, for the single-type model in gray. See table S1 for species name abbreviations.

| Model | Parameter | Posterior mean | Lower | Upper |
| --- | --- | --- | --- | --- |
| Single-type | $\lambda$ | 0.13 | 0.13 | 0.14 |
| Single-type | $\mu$ | 0.11 | 0.10 | 0.12 |
| Single-type | $\zeta$ | 5.44 | 3.81 | 7.51 |
| Single-type (fixed $\zeta$) | $\lambda$ | 0.13 | 0.13 | 0.14 |
| Single-type (fixed $\zeta$) | $\mu$ | 0.11 | 0.10 | 0.12 |
| Single-type (non extinct) | $\lambda$ | 0.14 | 0.13 | 0.14 |
| Single-type (non extinct) | $\mu$ | 1.00 | 0.87 | 1.15 |
| Single-type (non extinct) | $\zeta$ | 3.50 | 2.54 | 4.59 |
| Single-type (non extinct fixed $\zeta$) | $\lambda$ | 0.14 | 0.13 | 0.14 |
| Single-type (non extinct fixed $\zeta$) | $\mu$ | 1.01 | 0.88 | 1.15 |

**Figure S7:**
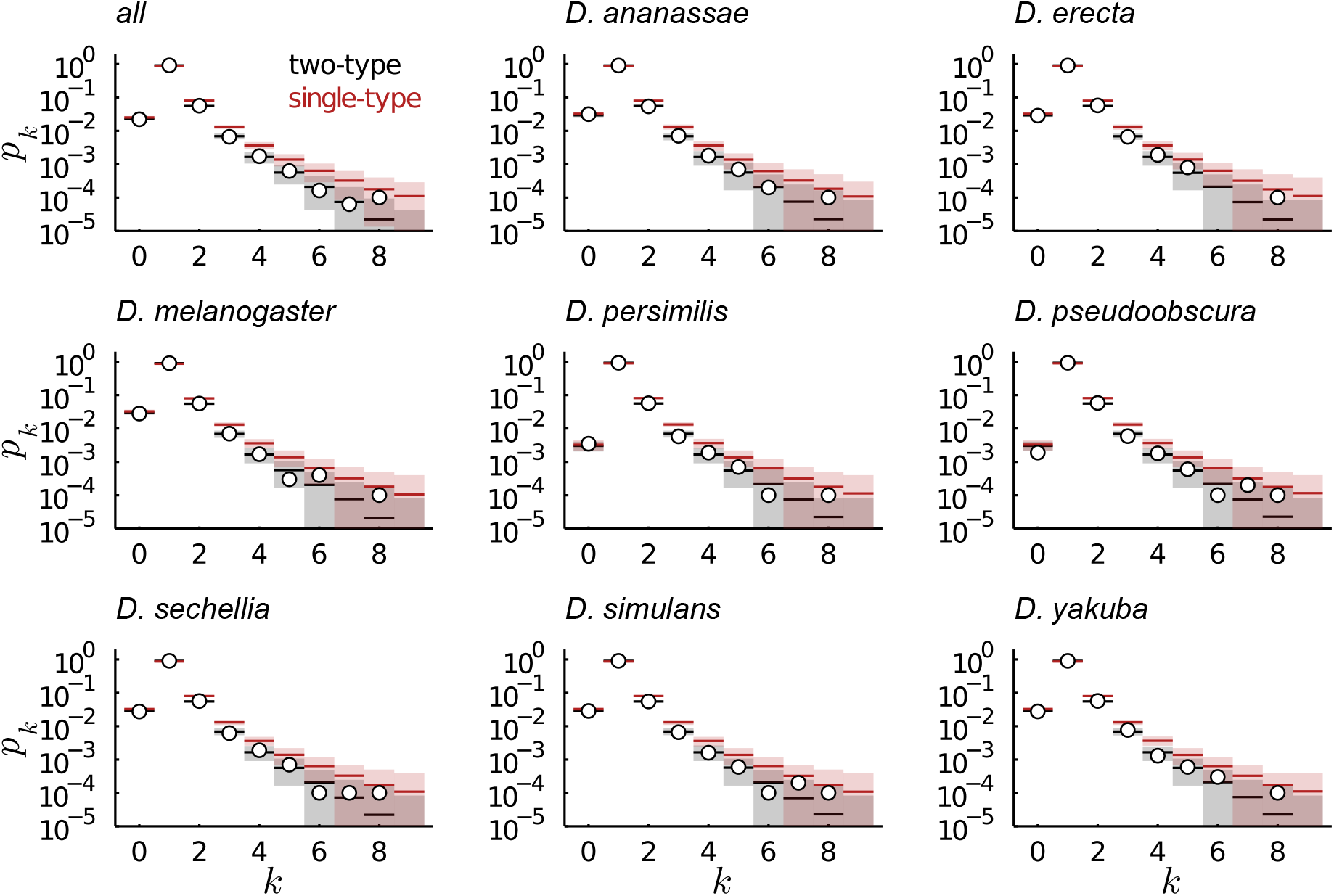
Posterior predictive simulations for the two-type DL model and the default single-type linear BDP model applied to data simulated under the DLF model.

**Figure S8:**
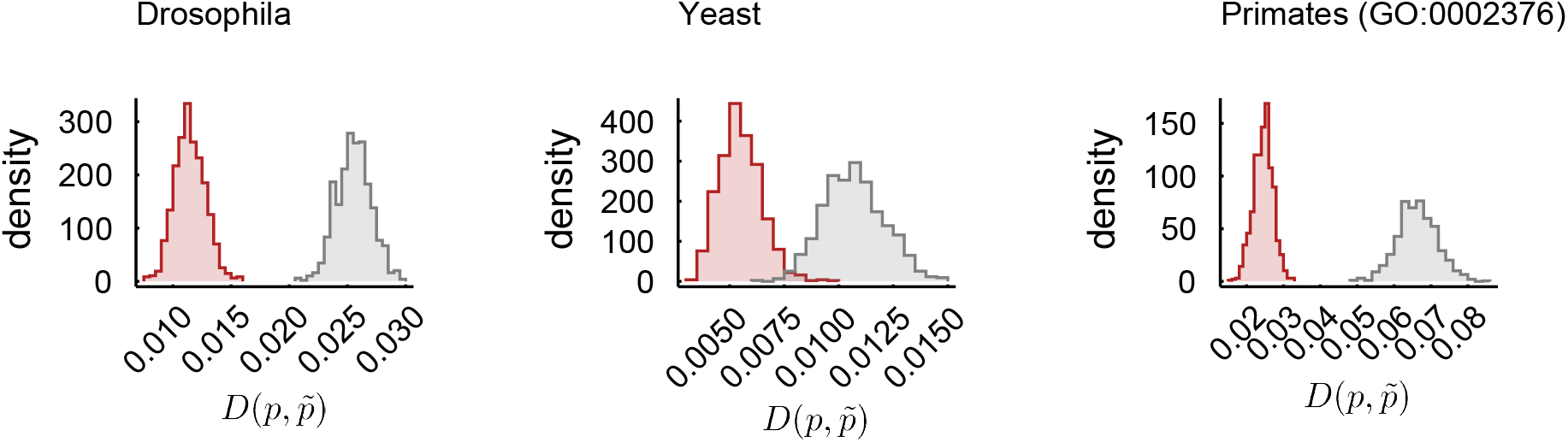
Kullback-Leibler (KL) divergence from posterior predictive simulations of the gene family size distribution to the observed family size distribution (across all species) for the *Drosophila*, yeast and primates data. In red the KL divergences for the simulations under the two-type model are shown, whereas in gray the distribution of divergences for the single-type model is shown.

**Figure S9:**
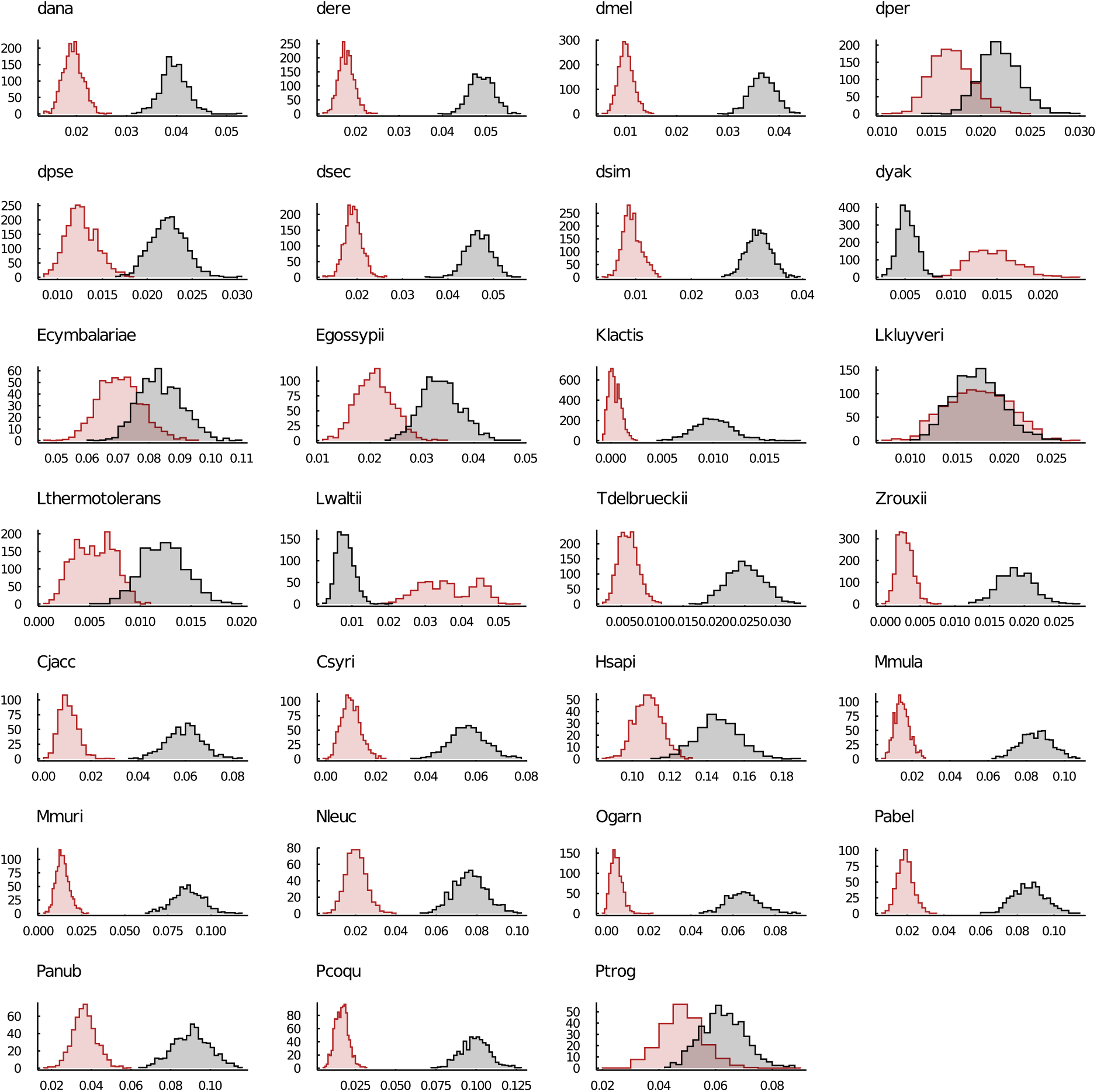
Kullback-Leibler (KL) divergence (along the *x*-axis) from posterior predictive simulations of the gene family size distribution to the observed family size distribution as in fig. S8, but now for each species separately. Results for the two-type model are displayed in red, for the single-type model in gray. See table S1 for species name abbreviations.

**Figure S10:**
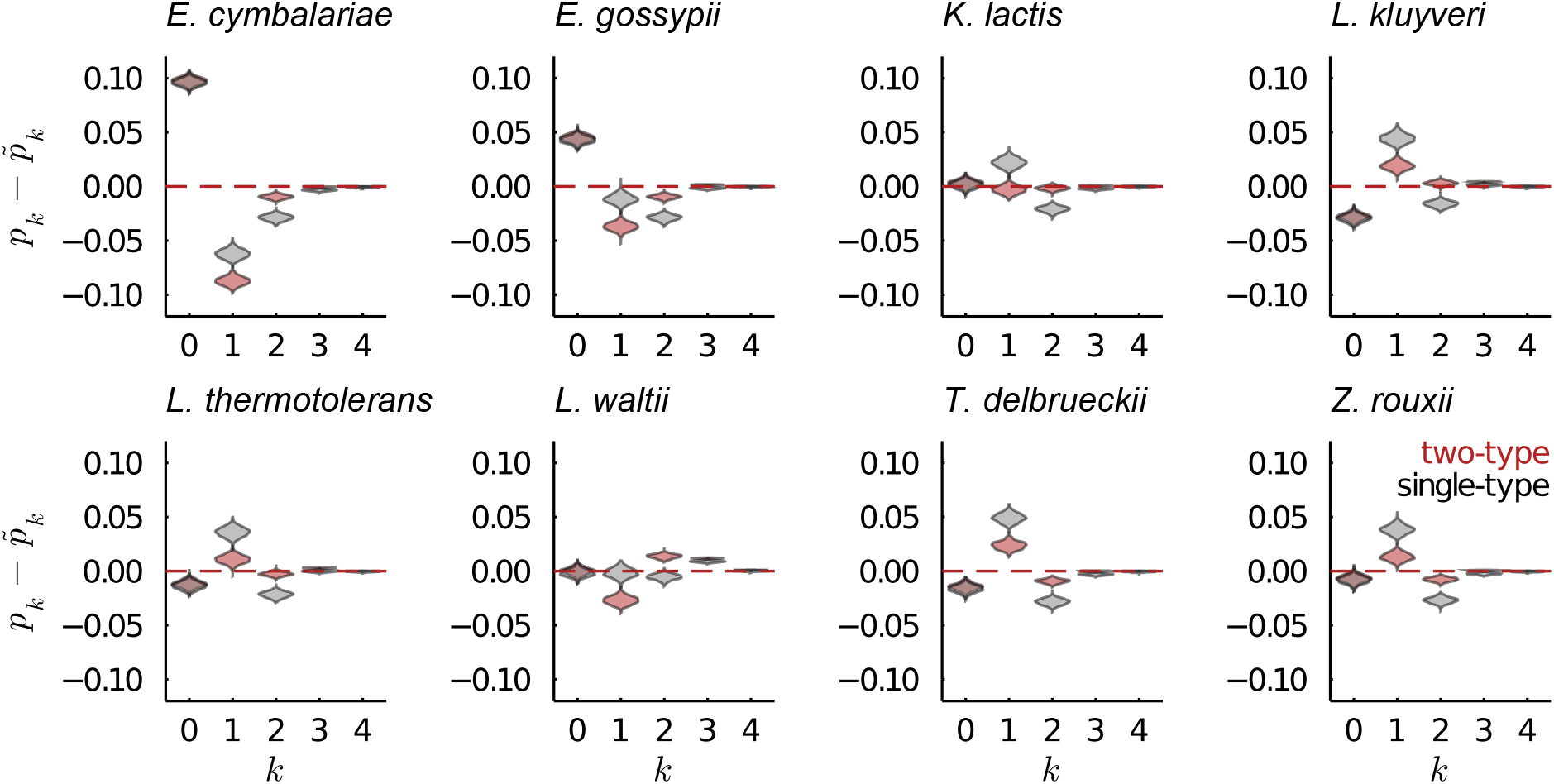
As fig. 4 but for the yeast data set.

**Figure S11:**
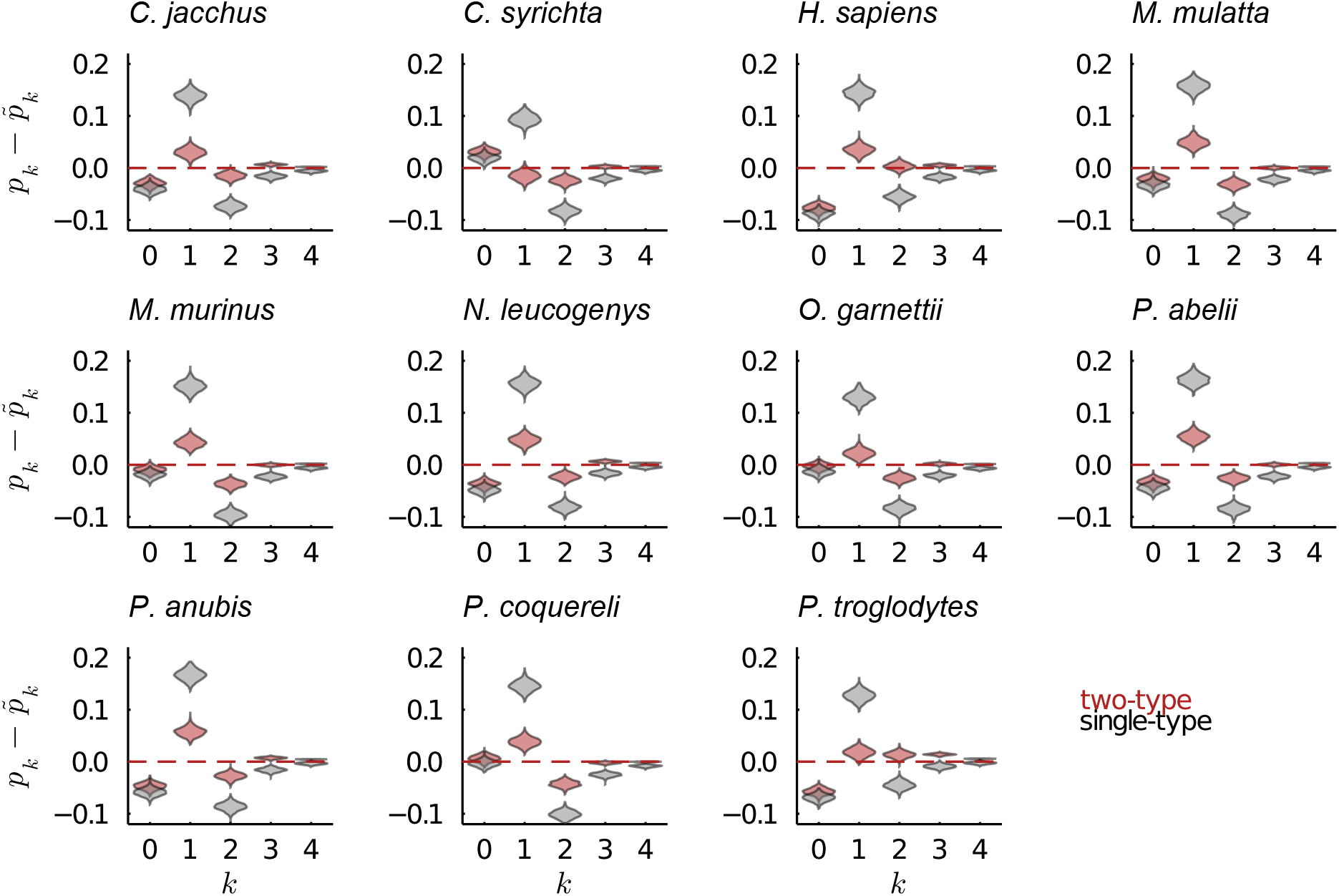
As fig. 4 but for the primates data set.

**Figure S12:**
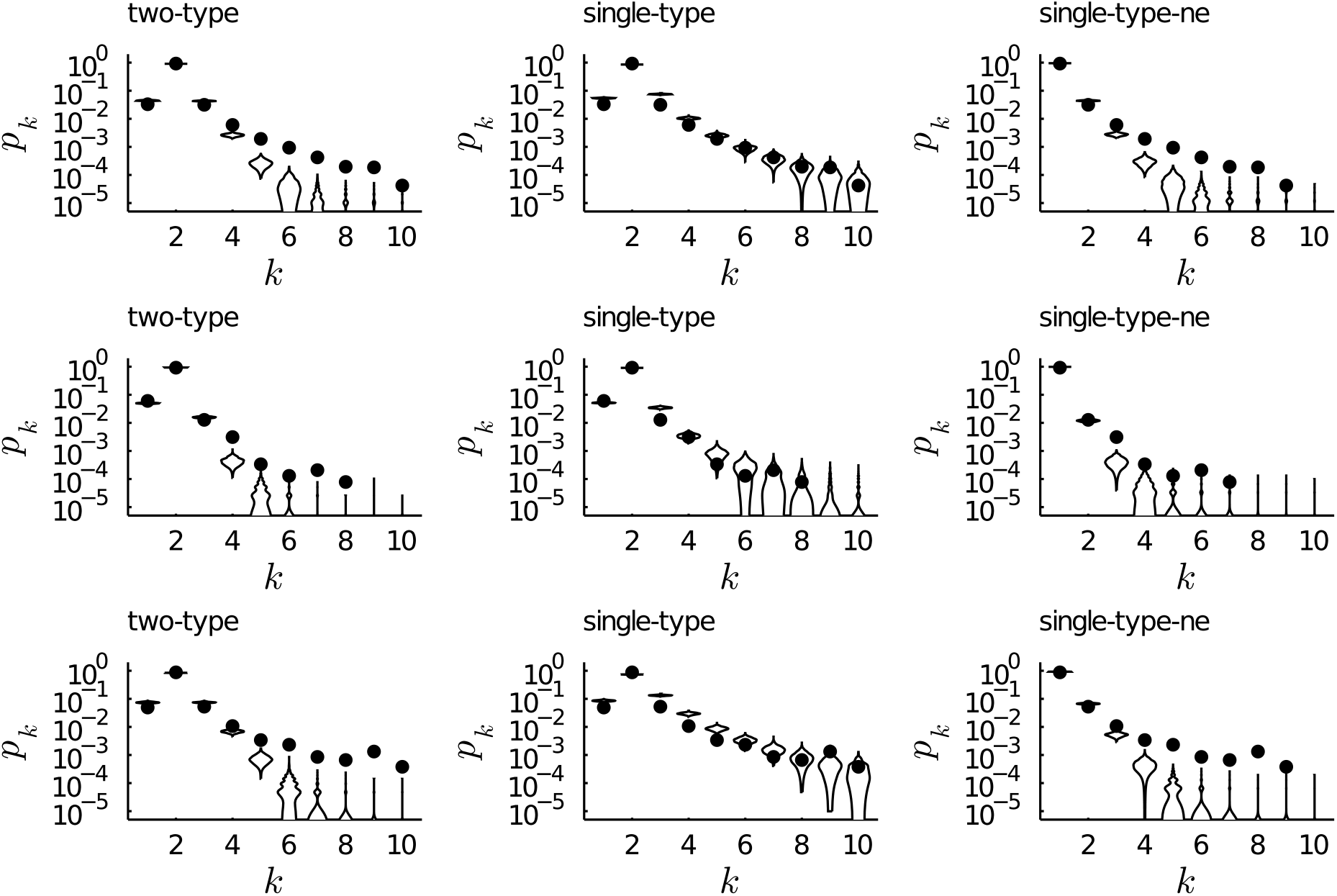
Posterior predictive family size distribution (across all lineages) for the *Drosophila* (top row), yeast (middle row) and primates data (bottom row) for the two-type, default single-type and single-type for non-extinct (‘single-type-ne’) families models. The dots show the observed size distribution.

